# Distinct functional classes of CA1 hippocampal interneurons are modulated by cerebellar stimulation in a coordinated manner

**DOI:** 10.1101/2024.05.14.594213

**Authors:** Jessica M. Froula, Jarrett J Rose, Chris Krook-Magnuson, Esther Krook-Magnuson

## Abstract

There is mounting evidence that the cerebellum impacts hippocampal functioning, but the impact of the cerebellum on hippocampal interneurons remains obscure. Using miniscopes in freely behaving animals, we find optogenetic stimulation of Purkinje cells alters the calcium activity of a large percentage of CA1 interneurons. This includes both increases and decreases in activity. Remarkably, this bidirectional impact occurs in a coordinated fashion, in line with interneurons’ functional properties. Specifically, CA1 interneurons activated by cerebellar stimulation are commonly locomotion-active, while those inhibited by cerebellar stimulation are commonly rest-active interneurons. We additionally find that subsets of CA1 interneurons show altered activity during object investigations, suggesting a role in the processing of objects in space. Importantly, these neurons also show coordinated modulation by cerebellar stimulation: CA1 interneurons that are activated by cerebellar stimulation are more likely to be activated, rather than inhibited, during object investigations, while interneurons that show decreased activity during cerebellar stimulation show the opposite profile. Therefore, CA1 interneurons play a role in object processing *and* in cerebellar impacts on the hippocampus, providing insight into previously noted altered CA1 processing of objects in space with cerebellar stimulation. We examined two different stimulation locations (IV/V Vermis; Simplex) and two different stimulation approaches (7Hz or a single 1s light pulse) – in all cases, the cerebellum induces similar coordinated CA1 interneuron changes congruent with an explorative state. Overall, our data show that the cerebellum impacts CA1 interneurons in a bidirectional and coordinated fashion, positioning them to play an important role in cerebello-hippocampal communication.

**Significance Statement:** Acute manipulation of the cerebellum can affect the activity of cells in CA1, and perturbing normal cerebellar functioning can affect hippocampal-dependent spatial processing, including the processing of objects in space. Despite the importance of interneurons on the local hippocampal circuit, it was unknown how cerebellar activation impacts CA1 inhibitory neurons. We find that stimulating the cerebellum robustly affects multiple populations of CA1 interneurons in a bidirectional, coordinated manner, according to their functional profiles during behavior, including locomotion and object investigations. Our work also provides support for a role of CA1 interneurons in spatial processing of objects, with populations of interneurons showing altered activity during object investigations.

## Introduction

Hippocampal spatial processing is shaped by both local and extrahippocampal elements. Within the hippocampus, this circuitry includes inhibitory interneurons (Freund and Buzsaki, 1996), which show a rich diversity of distinct activity profiles during different brain states and behaviors (e.g. (McNaughton et al., 1983; Colom and Bland, 1987; Mizumori et al., 1990; Katona et al., 2014, 2017; Lee et al., 2014; Varga et al., 2014; Arriaga and Han, 2017; Francavilla et al., 2018; Bocchio et al., 2020; Dudok et al., 2021a, 2021b; Szabo et al., 2022; Hainmueller et al., 2024). Hippocampal interneurons can profoundly impact local circuit function, including the formation of CA1 place fields (Grienberger et al., 2017; Dudok et al., 2021b). Indeed, some CA1 interneurons can even show spatial tuning themselves, although this has been noted to be significantly weaker than what is seen in CA1 pyramidal cells, (Kubie et al., 1990; Marshall et al., 2002; Maurer et al., 2006; Ego-Stengel and Wilson, 2007; Wilent and Nitz, 2007; Hangya et al., 2010; Geiller et al., 2020). Overall, much less is known about how hippocampal interneurons are involved in spatial information processing compared to principal cells. For example, CA1 pyramidal cells (especially deep CA1 pyramidal cells) often show landmark vector properties related to objects (Battaglia et al., 2004; Deshmukh and Knierim, 2013; Geiller et al., 2017); in contrast, little is known regarding how CA1 interneurons respond to objects (Tamboli et al., 2024).

Both chronic and acute manipulations of the cerebellum can also influence hippocampal spatial processing, including expression of CA1 place cells. For example, mice with impaired potentiation of Purkinje cells (i.e., “L7-PP2B” mice) appear to show an over-reliance on (faulty) self-motion information, with insufficient anchoring to local spatial cues, producing behavioral impairments on the Morris Water Maze and rotations of CA1 place fields (Lefort et al., 2019). Additionally, acute optogenetic manipulation of cerebellar Purkinje cells produces behavioral impairments on an object location memory task (but not an object recognition task) and reduces clustering of CA1 place fields near objects (Zeidler et al., 2020). Optogenetic manipulation of cerebellar Purkinje cells also produces acute altered firing of CA1 neurons (Choe et al., 2018; Zeidler et al., 2020). Previous work in our lab demonstrated that optogenetic activation of Purkinje cells in the vermis (lobule IV/V) or simplex produces an inhibition of activity of CA1 neurons during periods of light delivery, followed by rebound excitation immediately after light delivery (Zeidler et al., 2020). It is unclear if the inhibition of CA1 neuronal activity during Purkinje cell activation is due to external changes in inhibitory or excitatory inputs onto CA1 pyramidal cells, or if it is due to increased local inhibition.

Notably, prior work examining the impact of cerebellar manipulations on the hippocampus focused entirely (or effectively) on excitatory, principal, cell populations (Rochefort et al., 2011; Choe et al., 2018; Lefort et al., 2019; Zeidler et al., 2020). Thus, while prior work shows that the cerebellum can alter CA1 neuronal firing and, ultimately, functionality -- including objects-in-space processing -- it does not provide clear information on how cerebellar modulation impacts hippocampal inhibitory interneurons.

We therefore paired optogenetic excitation of Purkinje cells with concurrent miniscope calcium imaging of dorsal CA1 interneurons in freely moving animals to examine if cerebellar modulation impacts hippocampal interneurons. Given the dearth of information available regarding CA1 interneurons and the processing of objects in space, we additionally asked if any interneuron populations show increased (or decreased) activity related to objects and during object investigations, and if so, if these are in turn impacted by cerebellar modulation. More broadly, we examined how cerebellar modulation of interneurons corresponded with their functional activity profiles.

We find that optogenetic stimulation of cerebellar Purkinje cells modulates activity of a substantial portion of imaged CA1 interneurons. This includes both interneurons showing increased activity and interneurons showing decreased activity with cerebellar stimulation. These changes are not random, but instead occur in a coordinated manner across different interneuron populations and in line with their functional roles during behavior, including object investigations. Our findings illustrate a powerful impact of the cerebellum on hippocampal circuits, highlight a role of CA1 inhibitory interneurons in the processing of objects in space, and suggest that the cerebellum may influence processing in CA1 in part via bidirectional, coordinated, influences on populations of interneurons.

## Methods

All experimental protocols were approved by the University of Minnesota’s Institutional Animal Care and Use Committee.

### Animals

Both male and female mice were used in this study. Mice were bred in house and sexed at the time of weaning based on external genitalia. While this study was not powered to examine sex differences, we did not observe any large, overt, sex differences. To create mice selectively expressing channelrhodopsin (ChR) in cerebellar Purkinje cells, Pcp2-Cre mice (Jackson Laboratory #010536: B6.Cg-Tg(Pcp2-cre)3555Jdhu/J) (Zhang et al., 2004) were crossed with Ai32 mice (Jackson Laboratory #012569: B6; 129S-Gt(ROSA)26Sortm32.1(CAG-COP4*H134R/EYFP)Hze/J) (Madisen et al., 2012). Breeding pairs also generated non-opsin expressing animals, which were used as experimental controls. Prior to receiving head implants, mice were group housed. Following implantation, animals were individually housed to prevent damage to the implants. Animals were provided *ad libitum* access to food and water and were on a 14 h light/10 hr dark cycle.

### Surgical Procedures

All surgeries were performed stereotaxically while animals received 1%-3% isoflurane anesthesia and supplemental heating from a heating pad.

#### Viral injections

Mice were injected with AAV1-mDlx-GCaMP6f-Fishell-2 (titer: 4.43E13; developed in the lab of Gordon Fishell; purchased from Addgene: plasmid #83899, and packaged by the University of Minnesota Viral Vector and Cloning Core) to selectively label inhibitory interneurons in the hippocampus, with 500 nL of virus delivered at ∼100 nL per minute using a Hamilton Neuros Syringe (model 7002). Injection coordinates targeted the left dorsal hippocampal CA1 subregion (2 mm posterior, 1.75 mm left, and 1.5 mm ventral relative to bregma). After virus delivery, the syringe was left in place for 5 minutes before being retracted.

#### *Implants* (lens, optical fibers)

Following AAV1-mDLX-GCaMP6f transduction, a 1 mm circular craniotomy was made slightly off center from the viral injection site (specifically, centered at 2 mm posterior and 1.5 mm lateral to bregma). The cortex above the hippocampus was aspirated using a blunt needle (30-34 gauge) attached to a vacuum while ASCF was continuously applied. The aspiration was considered complete when axonal fibers of the alveus were visible and the field of view was cleared. Following aspiration, a gradient refractive index (GRIN) lens (Inscopix; 1mm wide, 4mm long) was lowered in place above CA1 using a stereotactic arm to a depth of 1.35 mm ventral relative to the skull at the posterior edge of the craniotomy. The lens was then covered with Kwik-Sil (World Precision Instruments) and an additional layer of cement to prevent damage. During the same surgery, animals were also implanted with optical fibers (RWD Life Science, 200µm core, 0.5 NA) in the cerebellum for light delivery to the cerebellar cortex. Fibers were implanted at a 10 degree angle (posterior from the vertical), targeting vermal lobule IV/V (6 mm posterior, 0 mm lateral, and at a depth of ∼0.01 mm with respect to the surface of the brain) and/or the right lateral simplex (5.7 mm posterior, 2.2 mm right, and at a depth of ∼0.01 mm with respect to the surface of the brain). Four animals were implanted with a fiber at only one location. Implants were secured to the head using cyanoacrylate, a small screw, and dental cement. Starting one day prior to surgery and for 3 days after surgery, animals received post-operative care of children’s motrin (10 mL/500 g water) in water for pain management. Animals also received a daily 100 μL injection of Dexamethasone (0.04 mg/mL in sterile saline) starting on the day of surgery and for 5 days post-op. At least 2 weeks after GRIN lens implantation, a baseplate was attached to the headcap with dental cement. During this procedure, the headcap was painted black to block any light leakage.

#### Posthoc verification

Following experiments, tissue was sectioned and examined to confirm correct placement of the GRIN lens and appropriate expression of mDLX-GCaMP virus. Post-mortem measurements of cerebellar optical fiber outputs yielded an average output of 1.7 ± 0.4 mW for implanted vermal fibers and 1.9 ± 0.3 mW for simplex fibers.

### Miniscope imaging and analysis

Calcium imaging data was acquired via a microendoscope (version 4.4 Miniscope; https://github.com/Aharoni-Lab/Miniscope-v4) (Cai et al., 2016) received pre-assembled from Open Ephys along with a custom DAQ PCB (Version 3.2; miniscope.org). This miniscope has a field of view up to 1 mm in diameter. Calcium videos were acquired by the Miniscope DAQ-QT software (https://github.com/Aharoni-Lab) at a rate of 20 frames per second with blue LED excitation (∼18-22% of max Miniscope output; ∼0.2-0.3 mW), gain (2-3.5x), and a focal plane set for each animal to yield a quality signal and field of view.

#### Processing of calcium signals

Calcium imaging videos were processed as described previously (Zeidler et al., 2020) using the MIN1PIPE calcium imaging signal extraction pipeline in Matlab 2021b ((Lu et al., 2018); https://github.com/JinghaoLu/MIN1PIPE). Briefly, MIN1PIPE employs strategies for background removal, movement correction, and automatic ROI identification/signal extraction. A priori structural element estimates were initialized at 5 pixels to seed ROI detection. From the resulting calcium transients, MIN1PIPE generates a ΔF/F signal and a deconvolved calcium signal, using a fast non-negative deconvolution method modified from (Pnevmatikakis et al., 2016).

### Light delivery

Animals received cerebellar light delivery once every two minutes via a DAQ (National Instruments) and custom TTL pulse generator software connected to a single channel LED driver and blue LED (Plexon; 465nm). A patch cable (Plexon, 200µm diameter, High Durability) to deliver light stimulations to either vermal lobule IV/V or the right lateral simplex was attached to the mouse for the duration of open field exploration (see details below in “Behavior” section). Stimulations were either 3 s of pulsed light delivery (frequency of ∼7Hz; 33% duty cycle: 50 ms on, 100 ms off) or a single pulse (1s duration). Delivery of stimulations occurred once every 2 minutes while the animal was in the arena, resulting in 25-30 stimulations per session per animal. During all recordings, a sleeve of tubing was used to cover the connection point between the implanted optical fiber and the patch cable to reduce light escape. Most animals (13/18) were tested with all combinations of stimulation frequency and site (i.e. 7Hz vermis, 7Hz simplex, 1s vermis, and 1s simplex), with at least 24 hours between each session. The remaining five animals were tested on a subset of stimulation paradigms either due to not having an optical fiber in both locations or due to technical difficulties. No attempt was made to track specific imaged cells across days of imaging.

#### Light responsiveness

The deconvolved calcium signal was used to analyze the responsiveness of individual hippocampal interneurons to cerebellar stimulation. The average calcium activity 3 seconds from the onset of stimulation was compared to the average calcium activity 3 seconds prior to the onset of stimulation across stimulation trials (25-30 per session) using a Wilcoxon Signed Rank test. We considered cells showing a significant response to light (p<0.05) to be light modulated (i.e., modulated by cerebellar manipulation). We additionally examined light responsiveness after correcting for locomotion-related impacts on calcium signals via linear regression **(Supplemental Figure 2, Figure 5)**. Determination of significant light modulation was otherwise similar (i.e. compared 3s periods using a Wilcoxon Signed Rank Test).

### Behavior

Prior to running animals in an arena setting, all mice were handled by an experimenter, habituated to the scope and patch cable attachments, and habituated to walking with the weight of the scope for at least 5 days.

#### Open field arena behavior

After scope and fiber attachment, each mouse had a 5-minute staging period in a small rectangular box before being moved into the main arena. The main arena was a large circular arena (∼50 cm diameter; wall height ∼30 cm) placed behind a curtain and gently illuminated via LED strip lights (∼100 lumens). During experiments examining the effects of light stimulation, no cues were present in the arena except an object in one quadrant. The object was filled with cement and was one of the following: a square glass votive holder (2”x2”x2”), a 50 mL glass beaker (2”Lx2”Wx3”H), or a circular tin (2.5”L x 2.5”W x 2”H). A sound machine placed under the center of the arena provided ∼80 dB white noise to reduce the influence of any outside sound. Animals were allowed to freely explore the arena for one hour while calcium data was recorded and cerebellar stimulations were delivered at regular intervals. Animal behavior footage was recorded by the Miniscope DAQ-QT software (concurrently collecting the calcium imaging data) at a rate of 15-30 frames per second from above the arena via a webcam (Logitech HD C270). Recorded behavior data was translated and aligned with respect to the calcium data timestamps for locomotion and object exploration analyses. Occasionally, tether detangling was necessary, and these periods of time were removed from further analysis. Between each animal, the arena and objects were wiped thoroughly with 70% ethanol and allowed to dry before the next animal’s recording.

#### Classification of behaviors

##### i) Locomotion

Each animal’s location in the arena was tracked using ezTrack ((Pennington et al., 2019); https://github.com/denisecailab/ezTrack) and then translated with respect to calcium video timestamps. Frame-by-frame X/Y coordinates were used to calculate speed and transitions between locomotion states (i.e. locomotion versus rest). Correlation of calcium signal with speed for the purposes of cell classification was assessed using only frames outside of periods of light delivery (i.e., for each light delivery, 30s of data, starting with the onset of light delivery, was excluded from this analysis). Slowdowns and speedups were defined by a point of transition between states, with a minimum of 0.5 seconds of walking followed by a minimum of 0.5 seconds of rest, or the reverse, respectively. An animal was deemed to be at rest if it moved less than 2.5cm in one second. Mean calcium activity on either side of a slowdown transition was compared to determine whether a cell showed increased activity during locomotion (“Loco↑” cell) or increased activity during rest (“Loco↓” cell). Similar results were found when instead using speedup transition states. Cells imaged during sessions with fewer than 15 instances of speedups or slowdowns were excluded from that particular classification.

##### ii) Object Exploration

Object investigations were manually scored by an experimenter blinded to experimental condition, using criteria as previously described (Vogel-Ciernia and Wood, 2014). Briefly, investigations included approach towards an object at close range (∼1 cm), sniffing the object, and rearing onto the object as long as the animal’s head was still oriented towards the object. A 4-second period immediately prior to the investigations was used as a paired baseline for analyses; investigations were excluded from comparisons if they had less than 4 s of non-investigation baseline. Mean calcium activity prior to onset of object investigations (3s, starting 4s prior to the onset of investigation) was compared to calcium activity during investigation (3s, starting with the onset of investigation) using a Wilcoxon Signed Rank test. The 1 s prior to object investigation was considered a “grace period” and excluded from the comparison to ensure baseline comparison reflected a true baseline. An Obj↑ cell was defined as a cell with a significant (p<0.05) increase in calcium signals during object investigations; Obj↓ cells showed a significant (p<0.05) decrease in calcium signals during object investigations. Cells imaged during sessions in which animals displayed less than 15 investigations were excluded from further object investigation-based analyses.

### Object responsiveness index analysis

Object responsiveness of CA1 interneurons was calculated using the location tracking data aligned to calcium activity (deconvolved signal) using an object responsiveness index (ORI) (Deshmukh and Knierim, 2011, 2013; Zeidler et al., 2020) and defined as ORI = (AO - AA)/(AO + AA), with AO being the mean rate of activity when the animal was within 7.5 cm of the object center and AA being the mean rate of activity when the animal was further than 7.5 cm from the object center. ORI values can range from -1 to 1. For each cell, its actual deconvolved data was compared to 2000 iterations of shifted location information. Cells were considered significantly responsive to objects if the ORI value was lower than 2.5% or higher than 97.5% of the shuffled ORI values (i.e., two-tailed test).

### Spatial information content analysis

Sessions in which the animal visited < 50% of the binned arena space were excluded from spatial information analysis. For our primary analyses, all data were speed filtered using an instantaneous speed threshold of 2.5 cm/s (i.e., only periods of locomotion were used). The location of the animal was binned into 0.5 cm square bins, then smoothed with a Gaussian kernel with sigma of 2.5cm to produce an occupancy map. Deconvolved calcium events were smoothed using the same Gaussian kernel and summed by bin. The smoothed neural activity was divided by the smoothed occupancy (by bin) to calculate the activity rate map for each cell. The spatial information content for each cell was defined as follows:

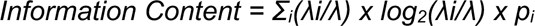

where λ is the mean calcium activity in the *i*-th spatial bin, and *p_i_*, is the probability of being in bin *i* (Skaggs et al., 1992; Rochefort et al., 2011; Shuman et al., 2020; Zeidler et al., 2020). A cell was deemed to have significant information content if its spatial information content score fell above the 95th percentile of 2,000 location-shuffled trials. A cell’s place field was defined as all bins with a smoothed activity rate of at least 50% of the activity rate of the bin with the greatest activity rate.

While a speed mask (i.e., limiting analysis of spiking information to only periods of time when the animal is locomoting) is typically used when examining pyramidal cell place cells, less is known about interneuron encoding of space and if this is locomotion-preferred. We therefore completed additional analyses, using all time points (not just periods of locomotion). However, we typically found stronger spatial information (SI) scores when restricting analyses to periods of locomotion (Supplemental Figure 3), and therefore used locomotion restricted data when calculating SI throughout.

### Statistics

Statistical analyses were conducted using OriginPro 2016 and MATLAB (version 2020b and 2023a). Data are presented as mean, with error bars and line shading represent SEM.

## Results

### Hippocampal interneurons respond to 7Hz stimulation of the cerebellar vermis

Previous research using calcium imaging to investigate the effects of cerebellar Purkinje cell stimulation on neurons in CA1 revealed a transient decrease in hippocampal neuron calcium activity during cerebellar stimulation, followed by large rebound in activity immediately following cerebellar stimulation (Zeidler et al., 2020). This previous work used a general neuronal promoter (hSyn). As excitatory pyramidal neurons make up ∼80% of the neurons in CA1 (Bezaire and Soltesz, 2013), this prior work primarily examined excitatory, pyramidal, neurons.

To investigate if cerebellar stimulation has an impact on GABAergic cells in CA1, we imaged CA1 interneurons while optogenetically activating cerebellar Purkinje cells. Specifically, in opsin-positive animals, the excitatory opsin channelrhodopsin (ChR2) was expressed selectively in Purkinje cells. Light was delivered via optic fiber to the cerebellar vermis (lobule IV/V; 3 seconds of ∼7Hz pulsed stimulation; 50 ms on, 100 ms off light pulses; matching previous work (Zeidler et al., 2020)). The calcium indicator GCaMP6f was expressed in dorsal CA1 interneurons using the mDLX enhancer (AAV-mDLX-GCaMP6f), and calcium activity was monitored via an attached miniscope in behaving animals freely exploring an open arena with a single object present.

Cerebellar stimulation in opsin-expressing animals modulated activity in a large proportion of imaged CA1 interneurons. Overall, approximately 43% (207 of 475 cells) of imaged neurons were significantly modulated by Purkinje cell activation **(Figure 1)**. CA1 interneurons responded heterogeneously to stimulation, with some cells showing increases in activity with light delivery (∼21%; hereafter referred to as Light↑ cells) and some showing decreases (∼22%; hereafter referred to as Light↓ cells) during cerebellar stimulation **(Fig. 1A-C)**. In contrast, opsin-negative animals exposed to the same experimental conditions showed only ∼5% of imaged interneurons as light-responsive, which is the expected false positive rate **(Fig. 1B-D)**. Accordingly, at the animal level, opsin-positive animals showed significantly more light-responsive interneurons, indicating that cerebellar modulation (rather than light delivery itself) produced the altered activity of CA1 interneurons (Mann Whitney U Test, p=0.0006; **Fig. 1D**). Findings were similar whether using the deconvolved signal (**Fig.1B-D**) or the ΔF/F signal (**Supplemental Figure 1**). We also performed additional analyses to control for any impacts of cerebellar modulation on locomotion of the animal and again found similar results **(Supplemental Figures 1 & 2)**, indicating that the influence of the cerebellum on the hippocampus is not mediated solely (or even predominantly) via indirect effects on locomotion.

**Figure 1.**
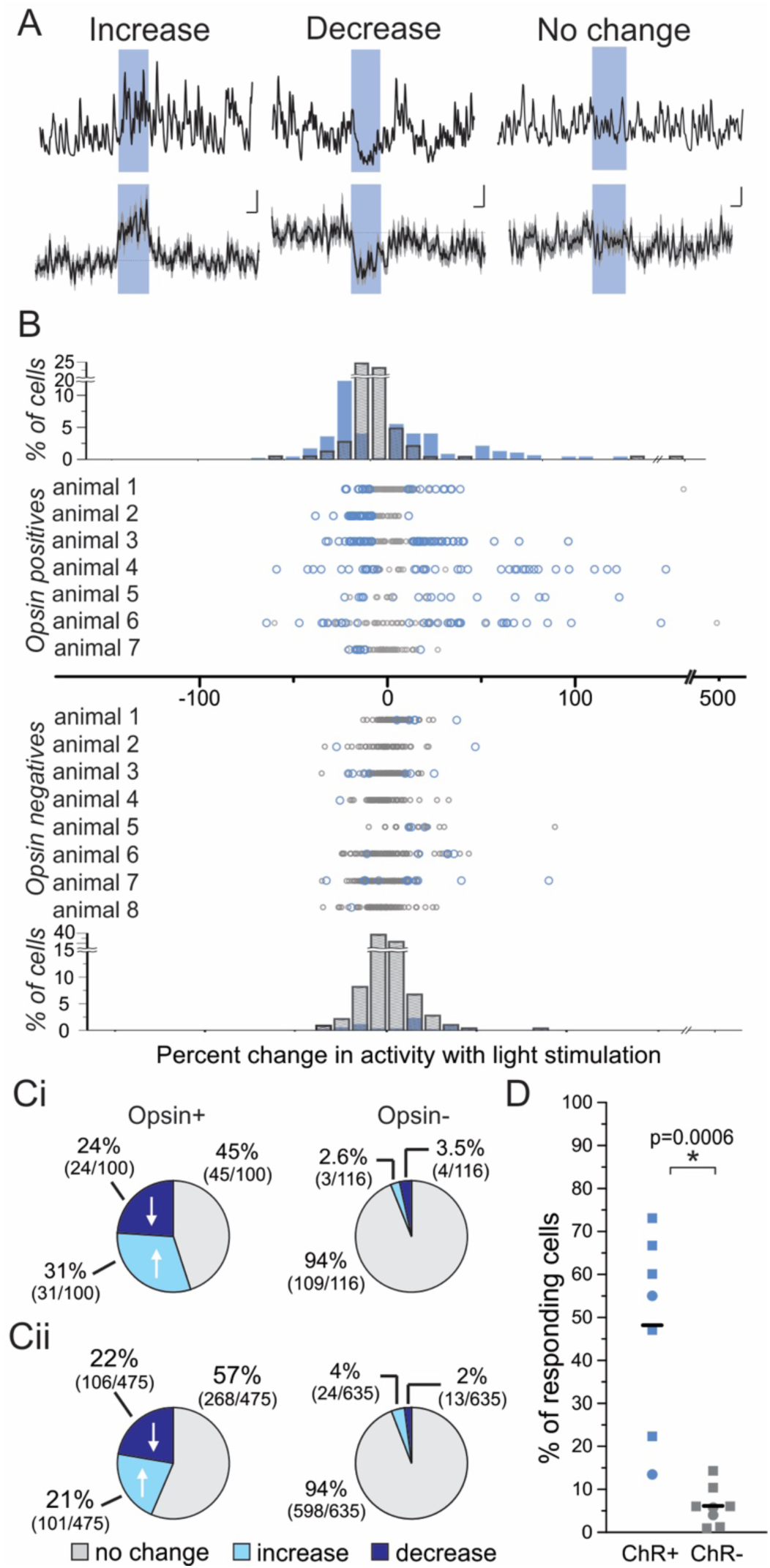
Cerebellar stimulation alters the activity of CA1 interneurons. **A,** Peristimulus calcium activity in example cells showing an increase (left), decrease (middle), or no change (right) with cerebellar stimulation. The stimulation period is represented by blue shading for all traces. Top: Example individual trace. Bottom: Average trace (black) for 25-30 stimulations in the same cell. Gray shading represents SEM. Scale bars: 1s, 0.05 ΔF/F (left, middle), 0.01 ΔF/F (right). **B,** Average percent change in interneuron activity with light stimulation. Each dot represents one cell. The histograms (top: opsin positives; bottom: opsin negatives) show the distribution of cells in each bin across all animals. Blue: cells significantly modulated by light delivery. Gray: cells not significantly modulated by light. **C,** Percentage and proportion of cells not responding to light (gray), showing a significant decrease in activity (dark blue), or showing a significant increase in activity (light blue), for an example opsin-positive and opsin-negative animal **(Ci)** and across all opsin-positive and opsin-negative animals **(Cii)**. **D,** Percent of responding cells by animal. Squares: males. Circles: females. Opsin-positive vs opsin-negative animals: p=0.0006, Mann-Whitney U Test.

Taken together, these data show that a large percentage of CA1 interneurons are responsive to acute cerebellar manipulation, indicating a previously uncharacterized role of hippocampal interneurons in cerebello-hippocampal interactions. CA1 interneurons do not respond in a homogenous manner to cerebellar stimulation, however, with some interneurons showing increases in activity and others showing decreases.

### Hippocampal interneurons are responsive to object exploration

We began analysis of interneurons’ spatial and object processing in opsin-negative animals, to avoid potential impacts of cerebellar modulation. As previously reported (Kubie et al., 1990; Marshall et al., 2002; Ego-Stengel and Wilson, 2007; Wilent and Nitz, 2007; Hangya et al., 2010), we found that CA1 interneurons’ firing could contain significant spatial information content (i.e., could be considered a “place cell”), but in part due to high background firing, CA1 interneurons typically show broad spatial tuning **(Figure 2A)**. However, we found a range of spatial information content scores and place field sizes **(Supplemental Figure 3)**. Some interneurons had more restricted spatial tuning profiles, including occasionally near the object **(Figure 2B)**. While the hippocampus is critical to the processing of objects in space, and pyramidal cells have been shown to change their firing in relation to objects (Komorowski et al., 2009; Manns and Eichenbaum, 2009; Burke et al., 2011; Deshmukh and Knierim, 2013; Gulli et al., 2020; Nagelhus et al., 2023), less is known about how interneurons respond to objects (Tamboli et al., 2024).

**Figure 2.**
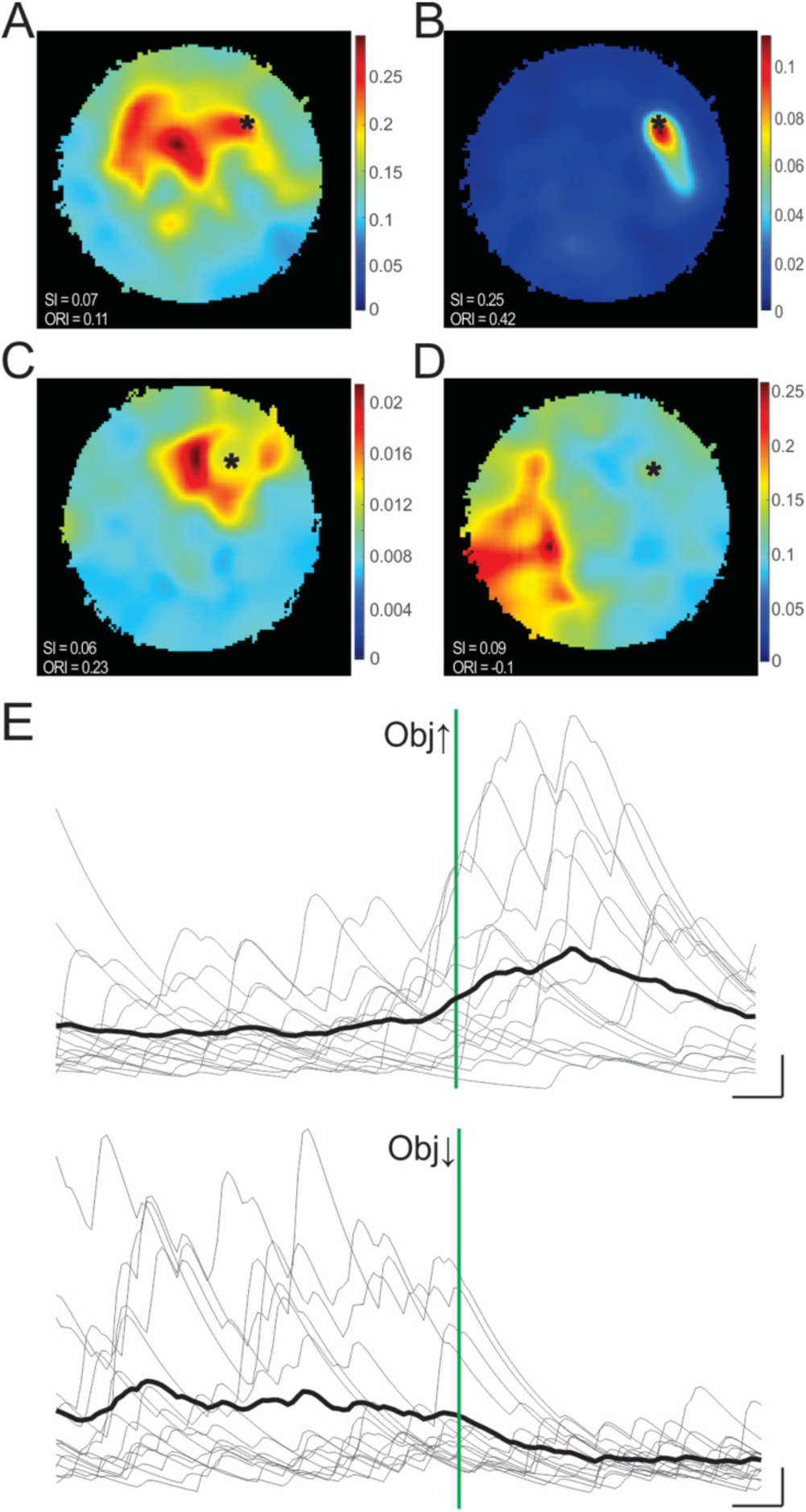
Hippocampal interneurons show modulated firing near objects and during object investigations. **A-D,** Example rate maps showing activity of CA1 interneurons during arena exploration with corresponding Spatial Information (SI) content scores and Object Responsiveness Index (ORI) scores. Black asterisk represents the location of the object in arena. **A,** An interneuron with broad spatial tuning. **B,** An interneuron with restricted spatial tuning near an object in the arena. **C,** An interneuron with a positive ORI. **D,** An interneuron with a negative ORI. **E,** Calcium activity aligned to the start of object investigations, in an example Obj↑ cell (top) and an example Obj↓ cell (bottom). Gray: individual traces. Black: Average ΔF/F. The green line indicates the start of the investigation. Scale bars: 0.5s; 0.1 ΔF/F (top), 0.02 ΔF/F (bottom).

We therefore examined the object responsiveness of interneurons using the object responsiveness index (ORI) (Deshmukh and Knierim, 2013; Zeidler et al., 2020), which compares cells’ activity near the object compared to activity elsewhere in the arena (**Figure 2C-D**). We found that some CA1 interneurons (21% of 498 cells in opsin-negative animals) showed preferential firing near objects (i.e., had a significant positive ORI value; **Fig. 2C**). This indicates that a substantial portion of interneurons have increased firing rates near objects. We also found that some CA1 interneurons had a significant negative ORI value (**Fig. 2D**), although these represented a smaller portion of imaged cells (9% of 498 cells). This suggests that other interneurons may actually show decreased activity due to objects. While this metric provides regional specificity, it only examines proximity to an object; it does not address whether or not the animal is actively exploring the object.

Therefore, we examined how interneuronal activity was specifically impacted by periods of object investigations, by comparing interneuron activity at the onset of object investigations to activity immediately prior (on average, 31 ± 3 object investigation periods were examined per animal). We found that a remarkably large percentage (∼30%) of imaged interneurons showed significantly altered activity during object investigations (183 of 635 interneurons in opsin-negative animals). The majority of investigation-modulated cells increased their activity with object investigation (hereafter referred to as “Obj↑” cells; 149 neurons) **(Figure 2E-F)**. A smaller proportion of interneurons decreased their activity during object investigation (hereafter referred to as “Obj↓” cells; 34 neurons). Cells’ activity profiles during object investigations could not be explained by changes in locomotor state **(Supplemental Figure 4)**. This data indicates that a substantial portion of CA1 interneurons show altered activity during object investigations, with some interneurons consistently showing an increase, and other interneurons consistently showing a decrease.

We were especially interested in interneuron processing of objects, as Zeidler et al. (2020) found that cerebellar stimulation impaired performance on an object location memory task and reduced hippocampal responsiveness to objects, with CA1 place cells mapping further away from objects (Zeidler et al., 2020). We therefore turned to see how cerebellar manipulation may impact interneuron spatial processing and object-responsiveness (**Figure 3**). As in opsin-negative animals, opsin positive animals (receiving periodic cerebellar stimulation) also had cells with significant spatial information (i.e. “place” cells), but the proportion of such cells was significantly reduced: 83% of imaged interneurons in opsin negatives vs 58% of imaged interneurons in opsin positive animals displayed significant information content (Opsin+ vs Opsin-, p=1.3×10^-20^, Chi-squared test, **Fig. 3A**). Overall, cells in opsin positives showed a reduced amount of spatial information (Opsin+ vs Opsin-SI content, p=6.4×10^-8^, Mann-Whitney U Test, **Fig. 3B**), with a trending but non-significant reduction of spatial information content in place cells specifically (Opsin+ vs Opsin-Place cell SI content, p=0.06, Mann-Whitney U Test). This suggests cerebellar stimulation is affecting spatial processing in CA1 interneurons.

**Figure 3.**
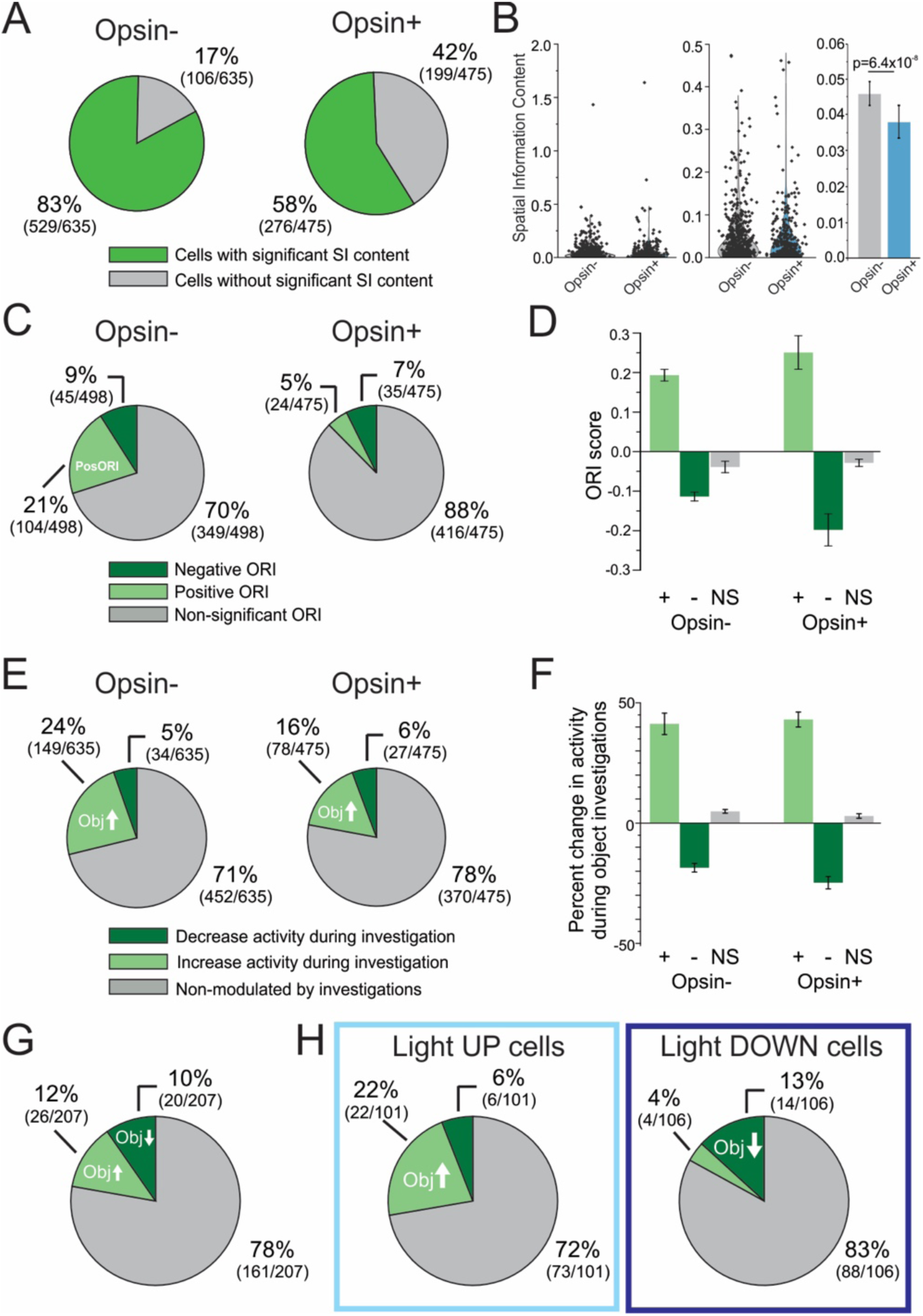
Cerebellar stimulation impacts spatial processing and object-responsiveness in CA1 interneurons. **A,** Proportion of imaged interneurons displaying significant spatial information (green) in opsin negative (left) and opsin positive (right) animals. Opsin positives have a reduced proportion of cells with significant SI content (Opsin- vs Opsin+, p=1.3×10^-20^, Chi-squared test), suggesting cerebellar manipulation impacts spatial information conveyed by CA1 interneurons. **B,** Left: Spatial information (SI) content scores across all cells in both genotypes. Note the large spread. Middle: distribution of SI content across cells, focused on smaller SI scores; Right: Average SI content in opsin negatives vs opsin positive animals. Interneurons in opsin-positive animals show smaller IC scores (p=6.4×10^-8^, Mann-Whitney), suggesting cerebellar stimulation impairs interneuronal spatial information content. **C,** Proportion of interneurons with a positive ORI (light green), a negative ORI (dark green), or a non-significant ORI (gray). Opsin positives have a reduced number of cells with a significant ORI value (Opsin- vs Opsin+; p=5.1×10^-13^; Chi-squared test), suggesting cerebellar stimulation reduces object responsiveness in CA1 interneurons. **D,** Average ORI scores by genotype with colors as in panel C. While there are fewer cells with significant ORIs in the opsin positive animals, for cells with a significant ORI, ORI scores are not significantly different between genotypes (Positive ORI scores in Opsin- vs Opsin+, uncorrected p=0.19, Mann Whitney; Negative ORI scores in Opsin- vs Opsin+, uncorrected p=0.15, Mann Whitney). **E**, Proportion of Obj↑ (light green), Obj↓ (dark green), and cells which did not show significant changes (gray) in activity during object investigations across genotypes. These distributions are significantly different (p=1.8×10^-4^, Chi-squared test), further indicating that cerebellar stimulation affects object-responsiveness in CA1 interneurons. **F,** Percent change in activity during object investigations as compared with baseline activity, with colors as in panel E to represent an Obj↑, Obj↓, or non-significantly responding cell. While fewer cells are object-responsive in opsin positive animals, in cells that are positively or negatively object responsive, the magnitude of that response is similar between genotypes (Obj↑ %change in Opsin- vs Opsin+, uncorrected p= 0.04, Mann Whitney; Obj↓ %change in Opsin- vs Opsin+, uncorrected p=0.03, Mann Whitney). **G,** The proportion of light-modulated neurons in opsin-positive animals that are Obj↑, Obj↓, or are not object-responsive. **H,** The proportion of Light↑ cells (left) or Light↓ cells (right) in opsin-expressing animals that are Obj↑, Obj↓, or are not object-responsive. Note that Light↑ cells that are object responsive are more likely to be Obj↑ cells than Obj↓ cells (left), while Light↓ cells that are object responsive are more likely to be Obj↓ cells rather than Obj↑ cells (right), suggesting coordinated modulation by cerebellar stimulation (Obj↑ vs Obj↓ in Light↑ and Light↓ cells, p=2.9×10^-5^, Chi-squared test). Numbers in parentheses for each pie chart section provide the number of cells. Data drawn from 7 opsin-positive animals (A-H) and 8 opsin negative animals (A-B, E-F) or 6 opsin negative animals (C-D).

Cerebellar stimulation also altered object responsiveness in CA1 interneurons. Opsin positive animals had a reduction in the proportions of cells with a significant ORI, with 12% of cells in opsin positive animals having either a positive or negative ORI compared to 30% of cells in opsin negative animals (Opsin+ vs Opsin-, p=5.1×10^-13^, Chi-squared test, **Fig. 3C**). The strength of ORI values in cells which showed a significant ORI were similar (**Fig. 3D**).

Similarly, interneurons were less likely to show altered activity during object investigations with cerebellar manipulation, and in particular, there were fewer Obj↑ cells: 24% of interneurons showed significant increases in activity during object explorations in opsin-negative animals, but only 16% did in opsin positive animals (proportions in Opsin+ vs Opsin-, p=1.8×10^-4^, Chi-squared test, **Fig. 3E**). This suggests that cerebellar stimulation may interfere with object processing in CA1 interneurons. The magnitude of firing rate changes during object investigations, in significantly modulated cells, remained similar (**Fig. 3F**).

Collectively, these data highlight that hippocampal interneurons are engaged during object processing, and that object responsiveness in CA1 interneurons can be impacted by cerebellar stimulation. In particular, the number of interneurons which are object responsive is decreased in animals receiving cerebellar stimulation.

### Bidirectional, coordinated, cerebellar modulation of object-related hippocampal interneurons

We next examined how interneurons that were responsive to cerebellar stimulation specifically responded to objects **(Fig. 3G)**. Of CA1 interneurons that were responsive to cerebellar stimulation, approximately 20% (46/207) also showed changes in activity during object investigations, indicating a partial overlap in light-responsive and object-responsive CA1 interneurons. Within this population of light-responsive interneurons that were also object responsive, there were interneurons which increased their activity during object investigations (Obj↑, 26/207 light-modulated cells) and interneurons which decreased their activity during object investigations (Obj↓, 20/207 light-modulated cells), suggesting that the cerebellum is able to impact both forms of object-responsive interneurons **(Fig. 3G)**.

When separated by direction of light modulation (increase or decrease), an interesting pattern emerged: cells that increased their activity with cerebellar stimulation were more likely to increase rather than decrease their activity during object explorations (22 Obj↑ vs 6 Obj↓ cells of 101 Light↑ cells), whereas cells that decreased their activity with cerebellar stimulation were more likely to decrease their activity during object explorations (14 Obj↓ vs 4 Obj↑ cells of 106 Light↓ cells) (Obj↑ vs Obj↓ in Light↑ and Light↓ cells, p=2.9×10^-5^, Chi squared test) **(Fig. 3H)**. This illustrates that the impacts of cerebellar modulation on hippocampal interneurons is not random, but rather is in accordance with their functional role – specifically, the direction of their modulation during object investigations. These data also help explain the heterogeneous response of hippocampal interneurons to cerebellar modulation **(Fig. 1).** Specifically, cerebellar modulation increases activity in subsets of Obj↑ CA1 interneurons and decreases activity in subsets of Obj↓ CA1 interneurons.

### Coordinated cerebellar modulation of functional interneuron populations extends to interneurons classified by locomotion activity profiles

Different hippocampal interneurons show different activity profiles related to locomotion (Lapray et al., 2012; Katona et al., 2014; Arriaga and Han, 2017; Hainmueller et al., 2024). For example, parvalbumin+ (PV) basket and axoaxonic cells, Type 3 interneuron selective interneurons (ISI), and early-born GABAergic ‘hub’ cells have been shown to be more active during movement (Bocchio et al., 2020; Luo et al., 2020; Dudok et al., 2021a, 2021b). In contrast, cholecystokinin+ (CCK) basket cells, M2R-expressing TORO (theta-off ripple-on) cells, and long-range projecting vasoactive intestinal peptide (VIP) cells have been shown to have higher activity levels during rest (Francavilla et al., 2018; Geiller et al., 2020; Dudok et al., 2021a; Szabo et al., 2022). Given the dichotomy we observed in cerebellar-responses with respect to object-responsiveness, we next asked if a similar dichotomy can be observed for interneurons functionally classified by their locomotion-activity profiles.

First, we examined the correlation of calcium activity (using the ΔF/F signal) to animals’ speed and identified cells whose activity was positively correlated, negatively correlated, or uncorrelated with locomotor speed **(Figure 4A-D)**. Note that this categorization was done using time periods outside of light delivery. As expected (Fox and Ranck, 1975; Colom and Bland, 1987; Mizumori et al., 1990), activity was correlated with speed in a majority of cells, including ∼97% of imaged cells (614/635) in opsin negatives and ∼95% of imaged cells (452/475) in opsin positives **(Fig. 4A)**, and the majority of these were positively correlated with speed **(Fig. 4B)**. There was a large range of correlations, and it should be noted that some significantly modulated cells were only weakly speed-modulated **(Fig. 4A, insets)**.

**Figure 4.**
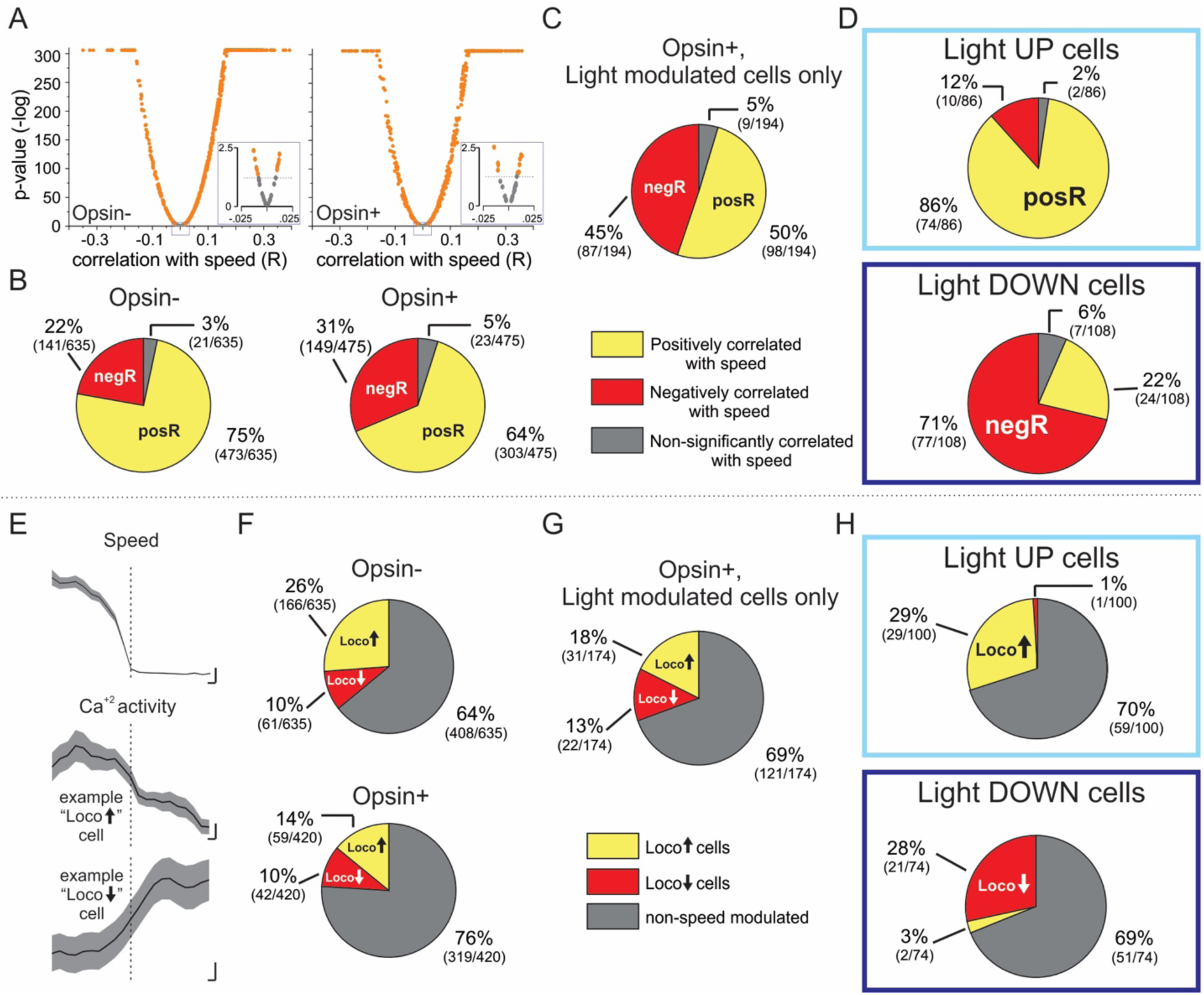
Cerebellar stimulation bidirectionally modulates CA1 interneurons in accordance with functional roles during locomotion. **A,** For each imaged interneuron, the correlation of animal speed and calcium activity, using the ΔF/F signal. y-axis: p-value (plotted on a negative log scale). Values below 10^-306^ are shown at p=10^-306^. x-axis: Spearman’s Rho (R). Left: data from opsin-negative animals. Right: data from opsin-positive animals. Each dot represents one cell. Significant correlation values (uncorrected p<0.05) are in orange and non-significant correlations are in gray. Insets: A zoomed-in portion of the graph, illustrating the cutoff point (p=0.05) between cells deemed significant and non-significantly speed-correlated. Note that many significantly correlated cells are only weakly (e.g. R<0.025) correlated. **B,** Proportions of positively correlated (yellow), negatively correlated (red), and non-speed correlated (gray) interneurons across genotypes. **C,** As in (B) but limited to only light-modulated cells in opsin-positive animals. **D,** The proportions of positively- and negatively-speed correlated cells for Light↑ (top) and Light↓ (bottom) interneurons in opsin-expressing animals. Note that most (86%) Light↑ interneurons that are speed-correlated are positively speed-correlated (top), while most (73%) Light↓ interneurons that are speed-correlated are negatively speed-correlated (bottom), suggested coordinated cerebellar modulation. **E,** Top: An example animal’s speed during 53 slowdowns. Middle: associated ΔF/F calcium activity in an example Loco↑ cell. Bottom: associated ΔF/F calcium activity in an example Loco↓ cell. Vertical dotted line represents the identified point of transition from movement to rest. For all, black: average; gray shading: SEM. Scale bars: 100 ms, 1 cm/s (top); 0.01 ΔF/F (middle, bottom). **F,** Proportions of Loco↑ (yellow), Loco↓ (red), and non-speed modulated (gray) cells by genotype. **G,** Similar to F, but for light-modulated cells in opsin-positive animals. **H,** Proportions of Light↑ cells (top) and Light↓ cells (bottom) in opsin-expressing animals that are Loco↑ (yellow), Loco↓ (red), or non-speed modulated (gray). Note that most Light↑ cells that are locomotion modulated are Loco↑ cells (top), while most Light↓ cells that are locomotion modulated are Loco↓ cells (bottom), again suggesting coordinated modulation by cerebellar stimulation. Data drawn from 7 opsin-positive animals (ΔF/F signal; A-D) or 6 opsin-positive animals (deconvolved signal; F-H; note that one animal did not have a sufficient number of slowdowns for inclusion in this analysis) and 8 opsin-negative animals (A,B,F).

When we specifically examined interneurons that were modulated by cerebellar stimulation, we again found a high percentage of neurons showing a correlation with speed, with 96% (185/194) of light-modulated interneurons showing a significant correlation with speed. Light-modulated cells were roughly equally split between being positively (50%) or negatively (45%) correlated with speed **(Fig. 4C)**. As with object-responsiveness, separating the light-modulated interneurons according to their functional properties as relates to locomotion yielded highly informative results **(Fig. 4D)**. Cells that increased their activity with cerebellar stimulation were cells that were primarily (86%) positively correlated with locomotion (74 positively correlated with speed, vs 10 negatively correlated with speed, of 86 Light↑ interneurons), and cells that decreased their activity with cerebellar stimulation were cells that were primarily (71%) negatively correlated with locomotion (77 negatively correlated with speed, vs 24 positively correlated with speed, of 108 Light↓ interneurons) **(Fig. 4D)**. This underscores that the cerebellum is able to modulate groups of interneurons in hippocampal CA1 in a coordinated manner, according to their function during behavior.

Given that this method broadly categorizes cells as positively or negatively correlated with speed, even if the relationship may not be particularly strong (i.e., a weak R value), we also examined the relationship of a cell’s activity to locomotion in an additional manner. Specifically, akin to previous work (Geiller et al., 2020; Dudok et al., 2021a), we identified periods of time that were transitions between movement states (e.g. a “slowdown”) **(Fig. 4E)**. Mean calcium activity on either side of this state transition was compared for each recorded cell to determine its classification as a cell that had higher activity during locomotion periods (hereafter referred to as “Loco↑” cells), a cell that had higher activity during rest periods (hereafter referred to as “Loco↓” cells), or a non-speed modulated cell. Using the slowdown categorization, fewer cells were classified as locomotion-modulated than using correlation values alone, but a substantial portion remained locomotion modulated **(Fig. 4F)**. Overall, a larger proportion were classified as being Loco↑ cells than Loco↓ cells: 26% were Loco↑ (166/635) vs 10% Loco↓ (61/635) in opsin-negative animals; 14% of cells were Loco↑ (59/475) vs 10% Loco↓ (42/475) in opsin-positives animals **(Fig. 4F)**.

When we specifically examined cells in opsin-positive animals that were responsive to cerebellar stimulation, we found that cerebellar modulation affected both Loco↑ cells (18%; 31 of 174 light-modulated cells) and Loco↓ cells (13%; 22 of 174 light-modulated cells) **(Fig. 4G)**. Importantly, among locomotion-modulated cells, we found that cells that increase their activity during cerebellar stimulation were overwhelmingly Loco↑ cells: 29% were Loco↑ cells (29 of 100 Light↑ interneurons) compared to only 1% Loco↓ cells (1 of 100 Light↑ cells). Conversely, cells that decrease their activity during cerebellar stimulation were overwhelmingly Loco↓ rather than Loco↑ cells: 28% were Loco↓ cells (21 of 74 Light↓ interneurons) vs only 3% for Loco↑ (2 of 74 Light↓ interneurons) (Loco↑ vs Loco↓ in Light↑ and Light↓ cells, p=7.3×10^-6^, Chi-squared) **(Fig. 4H)**. Similar results were found using speedup transitions instead of slowdown transitions (i.e., Light↑ cells were primarily Loco↑ cells; Light↓ cells were primarily Loco↓ cells; **Supplemental Figure 5**).

In summary, every method we used to look at cell activity during locomotion vs rest showed that cerebellar stimulation resulted in a similar profile: cells with activity positively modulated by speed are more likely to be excited by cerebellar stimulation while cells with activity negatively modulated by speed are more likely to show decreased activity during cerebellar stimulation. This, together with the object-investigation data, underscores that the cerebellum modulates groups of CA1 interneurons according to their functional roles. The heterogeneity of response profiles to cerebellar stimulation **(Fig. 1)** is not random, but instead illustrates coordinated, bidirectional modulation that corresponds to functional subclasses of CA1 GABAergic interneurons.

### Hippocampal interneurons respond to cerebellar stimulation at an additional frequency and site in cerebellar cortex

While we focused on 7Hz stimulation of the vermis, previous literature shows that other stimulation sites (e.g. the simplex) and other stimulation frequencies (indeed, even single light pulses) targeting the cerebellum can also have impacts on the hippocampus (Krook-Magnuson et al., 2014; Choe et al., 2018; Streng and Krook-Magnuson, 2020; Zeidler et al., 2020). We therefore additionally tested the impact of optogenetic activation of Purkinje cells targeting the simplex lobule, and the impact of a single 1 second light pulse (to either the simplex or the vermis). To account for any potential differences in locomotion across different stimulation paradigms **(Supplemental Figure 1)**, we compared locomotion-corrected data in opsin positive animals between these paradigms. Note that testing the impact of a single 1-second light pulse also allowed us to examine if stimulating at theta (with our ∼7Hz stimulation paradigm) was critical to the differential impacts noted on Loco↑ and Loco↓ cells, or if similar results could be achieved without inducing a hippocampal theta oscillation (Zeidler et al., 2020).

At the animal level, different stimulation paradigms elicited different percentages of responding cells (7Hz vs 1s, p=0.05; vermis vs simplex, p=0.03; interaction, p=0.65; Two-way Repeated Measures ANOVA, **Figure 5**), suggesting that hippocampal interneurons may respond more or less strongly depending on where and how the cerebellum is stimulated. Specifically, vermal stimulation produces a stronger response than simplex stimulation – a finding previously reported (Krook-Magnuson et al., 2014; Zeidler et al., 2020). Additionally, a single one-second continuous light delivery may produce slightly larger responses than pulsed 50 ms light. This is somewhat in keeping with previous work that indicated a 50 ms pulsed cerebellar stimulation is more effective at evoking a hippocampal response than a 10 ms pulsed stimulation (Zeidler et al., 2020), suggesting longer pulse lengths may produce greater effects (which is perhaps not a surprising finding). Similarly, when we analyzed stimulation paradigms at the cell level, there were significantly different proportions of responding cells, with 1s stimulation producing more responsive cells than 7Hz pulsed stimulation at both sites (Vermis: 7Hz vs 1s, p=1.5×10^-9^; Simplex 7Hz vs 1s, p=1.1×10^-8^, Chi-squared tests) and vermis stimulation producing more responding cells than simplex stimulation at either frequency (7Hz Vermis vs Simplex, p=1.8×10^-14^; 1s Vermis vs Simplex, p= 1.3×10^-5^, Chi-squared tests) **(Fig. 5B-C)**.

**Figure 5.**
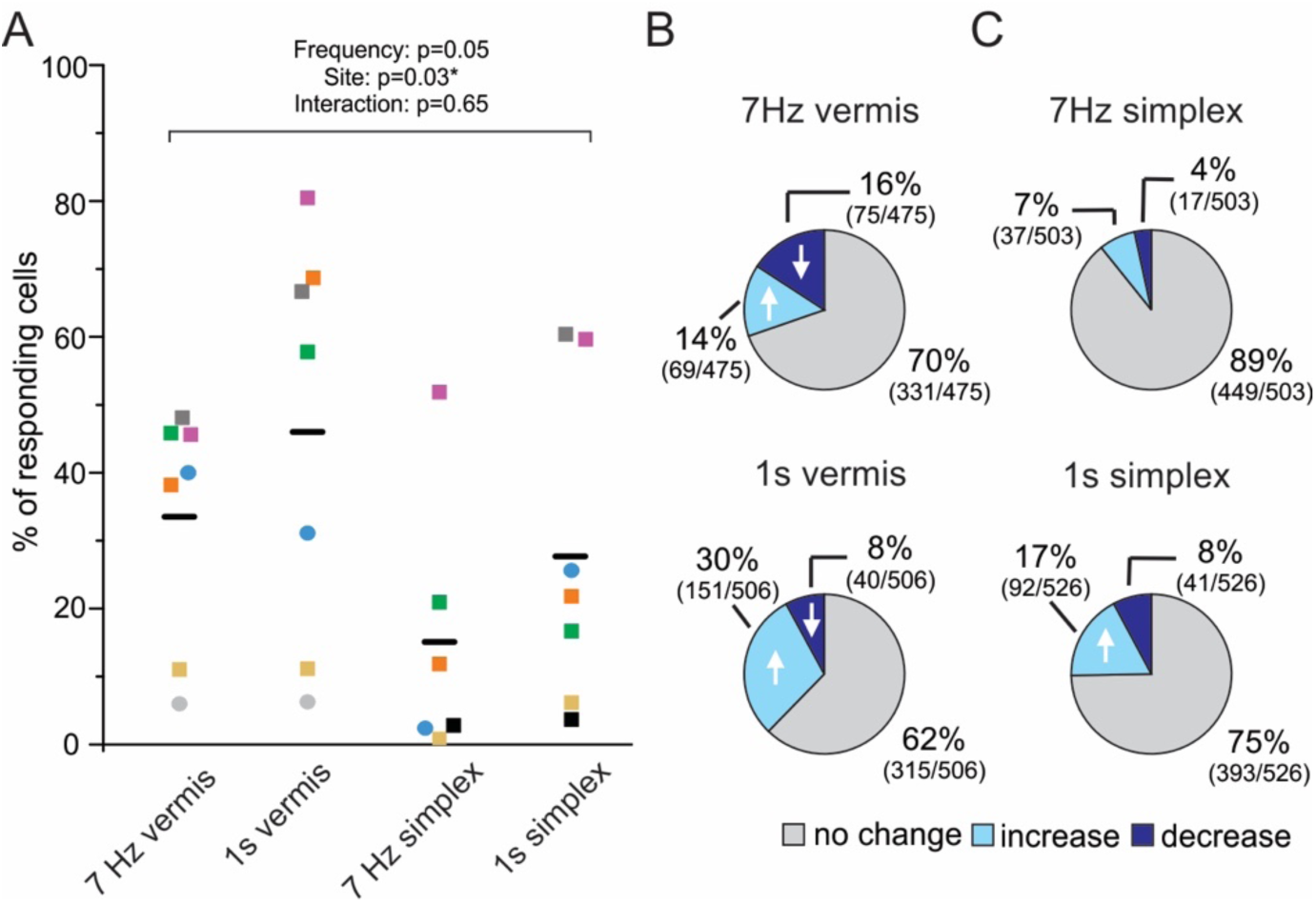
Hippocampal interneurons respond to cerebellar stimulation across light delivery paradigms. **A,** Percentage of light-responsive cells after locomotion correction across opsin-positive animals when stimulating the vermis or simplex at 7 Hz with 50ms light pulses or with a single, continuous, 1 second light pulse. There are slight differences between the percentage of responding cells across stimulation paradigms (7Hz vs 1s, p=0.05; vermis vs simplex, p=0.03; Interaction, p=0.65; Two-way Repeated Measures ANOVA. Only animals with data across all conditions (colored symbols) were included in the repeated measures ANOVA). Squares = males. Circles = females. **B,** Proportion of Light↑ (light blue), Light↓ (dark blue), and non-light modulated cells (gray) during vermis stimulation at 7 Hz or with a 1-second light pulse. **C,** Same as in (B), but for simplex stimulation. At the cell level, 1s stimulation was more effective than 7Hz pulsed stimulation at both sites (Vermis: 7Hz vs 1s, p=1.5×10^-9^; Simplex 7Hz vs 1s, p=1.1×10^-8^, Chi-squared tests). Additionally, vermis stimulation was more effective than simplex stimulation at either frequency (7Hz Vermis vs Simplex, p=1.8×10^-14^; 1s Vermis vs Simplex, p= 1.3×10^-5^, Chi-squared tests). Data drawn from 7 animals with vermis stimulation, 7 animals with 1 second simplex stimulation, and 6 animals with 7 Hz simplex stimulation.

We next examined how different stimulation paradigms modulated functionally defined populations of hippocampal interneurons (**Figure 6**). We found that the general profile of modulation with respect to object responsiveness held across stimulation paradigms, with Light↑ interneurons more likely to be Obj↑ rather than Obj↓ cells, and Light↓ interneurons being conversely more likely to be Obj↓ than Obj↑ cells **(Fig. 6A-D)**. With respect to locomotion, speed-modulated Light↑ cells were primarily Loco↑ cells and speed-modulated Light↓ cells were primarily Loco↓ cells, regardless of the cerebellar stimulation site (vermis or simplex) or paradigm (7Hz pulsed or 1 second continuous) (**Fig. 6E-H**).

**Figure 6.**
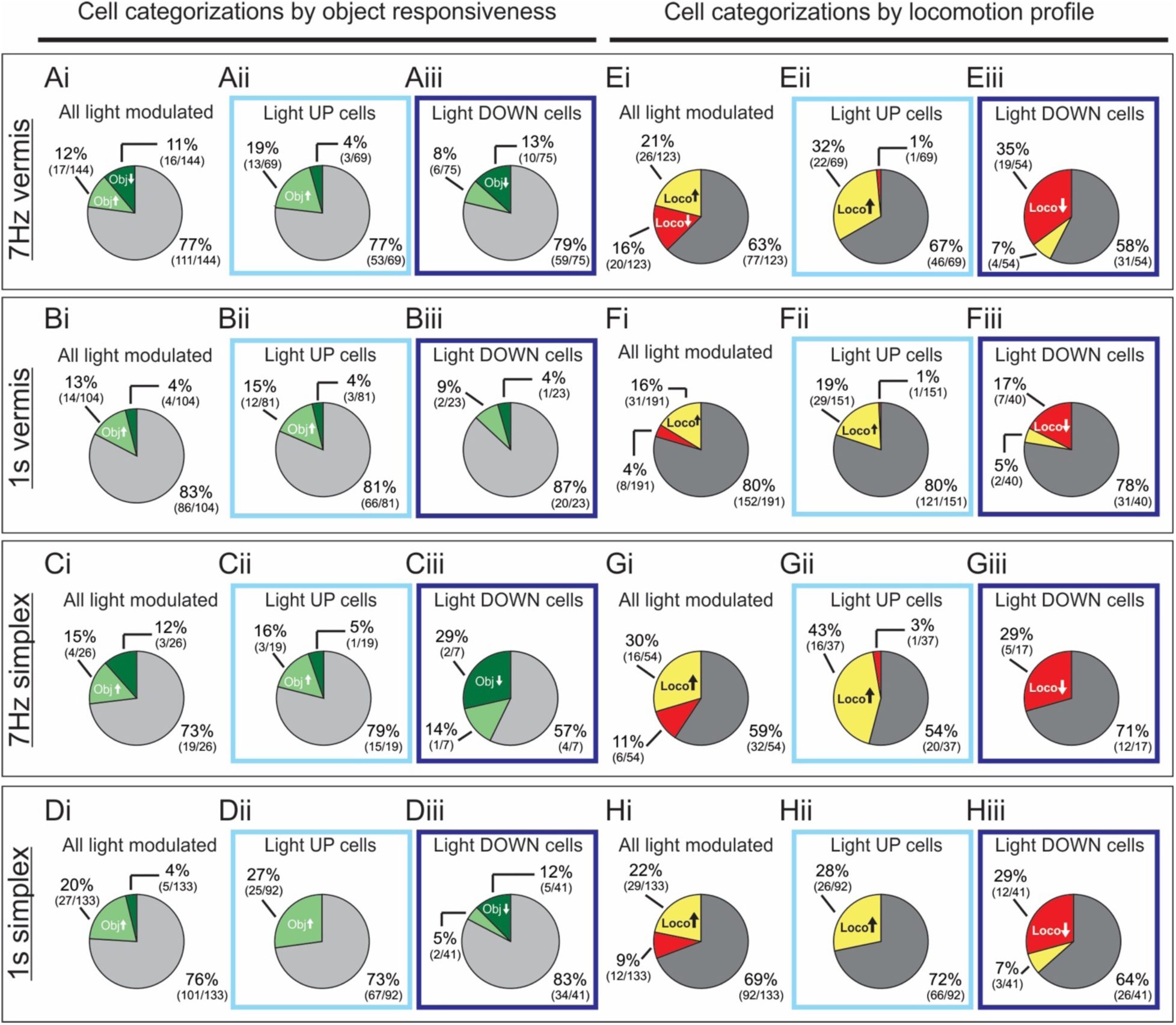
Coordinated cerebellar modulation of hippocampal interneurons occurs across stimulation paradigms. Light-modulated cells after locomotion correction in opsin-positive animals with stimulation of the cerebellum either in the vermis **(A,B,E,F)** or right lateral simplex **(C,D,G,H)** with either a 7Hz pulsed stimulation **(A,C,E,G)** or a 1 second single pulse stimulation **(B,D,F,H)**. Data either represent all light modulated cells (**i**) or are further parsed by light modulation profiles: Light↑ cells (**ii**, light blue boxes) or Light↓ cells (**iii**, dark blue boxes). Cell response profiles according to object responsiveness are on the left side (**A-D**), with light green representing Obj↑ cells, dark green representing Obj↓ cells, and gray representing cells that are not object responsive. Cell slowdown response profiles are on the right side (**E-H**), with Loco↑ cells in yellow, Loco↓ cells in red, and non-speed modulated cells in gray. Note that in general, across stimulation paradigms, Light↑ cells tend to be Obj↑ cells and Loco↑ cells rather than Obj↓ and Loco↓ cells, while Light↓ cells are more likely to be Obj↓ and Loco↓ cells. Statistical comparisons between 7Hz and 1s data can be found in **Supplemental Table 1**. Data is drawn from 7Hz vermis: 7 opsin-positive animals (locomotion and object data); 1s vermis: 7 (locomotion) and 4 (object) opsin-positive animals; 7Hz simplex: 6 (locomotion) and 5 (object) opsin-positive animals; 1s simplex: 7 (locomotion and object) opsin-positive animals.

Taken together, these results underscore that cerebellar (vermis or simplex) modulation bidirectionally impacts CA1 interneurons according to their functional profiles during behavior and indicates that this coordinated modulation is not dependent on the specific stimulation frequency used.

## Discussion

We show that cerebellar stimulation impacts the activity of a large portion of CA1 interneurons. These impacts are diverse -- with cerebellar stimulation evoking increases in some cells and decreases in others -- but not random. Rather, the cerebellum modulates CA1 interneurons in a coordinated manner, with changes reflecting their functional properties during behavior. Specifically, cerebellar stimulation increases activity in interneurons that show an increase in activity during locomotion or object investigations, and cerebellar stimulation concurrently decreases activity in other CA1 interneurons that show decreased activity during locomotion or object investigations. Our results also provide potential insight into how the cerebellum influences hippocampal processing of objects in space: we find that a subset of CA1 interneurons change their firing rate during object investigations, and that a subset of these cells are modulated by cerebellar stimulation. Collectively, our findings indicate diverse, but structured, influences of the cerebellum on the hippocampus, which may explain impacts on spatial processing, including the processing of objects in space.

Hippocampal spatial processing has repeatedly been shown to be influenced by cerebellar function (recently reviewed in (Froula et al., 2023)). For example, transgenic animals with disrupted long-term depression (LTD) of excitatory synapses from parallel fibers onto cerebellar Purkinje cells show intact use of spatial cues, but impaired use of self-motion cues, and unstable CA1 place fields in the dark (Rochefort et al., 2011). Conversely, transgenic animals with impaired long-term potentiation (LTP) in Purkinje cells show impaired orientation and unstable CA1 representations relative to a visual cue, suggesting the cerebellum is important for the processing of spatial cues or landmarks (Lefort et al., 2019). Notably, a subset of CA1 pyramidal cells, especially those in deep CA1 (Geiller et al., 2017), are influenced by the presence of objects (Komorowski et al., 2009; Manns and Eichenbaum, 2009; Burke et al., 2011; Deshmukh and Knierim, 2013; Gulli et al., 2020; Nagelhus et al., 2023), and this is disrupted by acute cerebellar manipulation. Specifically, in cerebellum-stimulated animals, cells in CA1 overall are less responsive to objects in an environment, and CA1 place fields map further away from objects (Zeidler et al., 2020). This change in CA1 firing properties is paired with behavioral deficits in processing objects in space: animals with cerebellar stimulation show impaired performance on an object location memory task (Zeidler et al., 2020). A similar deficit is not seen in a similarly formatted object recognition task, highlighting a critical role of cerebellar-hippocampal interactions in the processing of objects in space, rather than a more general processing of objects. Importantly, our current results highlight a possible role of interneurons in hippocampal processing of objects in space and show a cerebellar influence on CA1 interneurons’ processing of objects.

While a growing number of studies have examined the impact of objects on CA1 place cells, much less is known about the impact of objects on CA1 interneurons. This is especially important to understand, as hippocampal interneurons are likely critical for object-location processing. Broad activation of CA1 interneurons (Yu et al., 2018) or inactivation of SOM+ interneurons specifically (Honoré and Lacaille, 2022) (but not activation of PV interneurons (Zeidler et al., 2020)) impairs performance on a hippocampal-dependent Object Location Memory (OLM) task. We report that a significant portion of CA1 interneurons (∼30%) are responsive during object investigations, including both cells with increased and cells with decreased activation. These changes in activity during object investigations could not be explained by changes in locomotion **(Supplemental Fig. 4)**, but could potentially reflect specific exploratory behaviors, such as whisking or sniffing (Moore et al., 2013; Dudok et al., 2021b). While future work will be necessary to further interrogate activity patterns of interneurons related to object investigations, it is clear that hippocampal interneurons are critical to successful processing of objects in space (Yu et al., 2018; Honoré and Lacaille, 2022), and that hippocampal interneurons show altered activity during object investigations **(Fig. 2)**.

Importantly, we found that a subset of interneurons that were object responsive were also responsive to cerebellar manipulation. This provides a potential route by which the cerebellum can influence hippocampal processing of objects in space – that is, by influencing object-responsive CA1 interneurons. Additionally, we found that the number of interneurons with significant information content and the number of interneurons responsive to objects was decreased in animals receiving cerebellar modulation, further underscoring the influence of the cerebellum on CA1 processing of objects in space via interneurons.

Not only were object-responsive CA1 interneurons able to be modulated by the cerebellum, but also, this cerebellar mediated modulation of object-responsive CA1 interneurons occurred in a highly coordinated manner. CA1 interneurons that increased their activity with light were more likely to show increased activity during object investigations (rather than decreased activity), while CA1 interneurons that *decreased* their activity with light were more like to show *decreased* activity during object investigations. That is, the cerebellum was able to dial up activity in certain interneurons and while dialing down activity in other interneurons, in line with their activity profiles during object investigations. Interestingly, one theory regarding the role of the cerebellum is coordination of activity across brain regions (Popa et al., 2013; McAfee et al., 2019, 2022; Lindeman et al., 2021). Our work suggests the cerebellum also has coordinated effects on neuronal populations within a brain region.

Cerebellar modulation of interneurons occurring in a coordinated manner was also evident when examining interneuron activity profiles as relates to locomotion. The majority of CA1 interneurons show significant modulation across locomotor states (Fox and Ranck, 1975; Colom and Bland, 1987; Mizumori et al., 1990; Arriaga and Han, 2017; Szabo et al., 2022), with neurogliaform/Ivy cells being a notable exception (Fuentealba et al., 2008; Lapray et al., 2012; Geiller et al., 2020). Of those modulated by locomotion, the majority are positively modulated (Lapray et al., 2012; Geiller et al., 2020; Dudok et al., 2021b; Hainmueller et al., 2024). However, certain populations, including CCK-basket cells, M2R-expressing TORO (theta-off ripple-on) cells, and long-range projecting VIP cells have been shown to have higher activity during rest (Francavilla et al., 2018; Geiller et al., 2020; Dudok et al., 2021a; Szabo et al., 2022). This diversity of interneuron activity profiles (no change, increased, decreased) as relates to locomotion suggest different, and at times dichotomous (Dudok et al., 2021a), roles for different interneuron populations in hippocampal processing. These dichotomous roles can be imposed by external inputs (e.g. the medial septum), differential receptor expression (e.g. AChR vs M2R expression), and internal interactions (e.g., inhibition of CCK-basket cells by PV-basket cells and vice versa (Karson et al., 2009; Dudok et al., 2021a)). Importantly, we find that both Loco↑ and Loco↓ cells can be impacted by cerebellar modulation, indicating that diverse hippocampal interneurons are impacted by cerebellar modulation. Moreover, we find that this cerebellar modulation of CA1 interneurons occurs in a coordinated fashion across interneuron populations, such that Light↑ cells are typically Loco↑ (rather than Loco↓) and Light↓ are typically Loco↓ (rather than Loco↑) cells. This underscores that the cerebellum does not modulate interneurons in a random manner, but instead does so in a coordinated, bidirectional, manner, with certain neuronal populations being activated during cerebellar modulation and others being inhibited. This was true whether looking at activity profiles with respect to locomotion or with respect to object investigations.

Notably, both locomotion and investigative behaviors are associated with hippocampal theta states (Bland, 1986; Buzsáki, 2002; Grion et al., 2016; Kleinfeld et al., 2016). Our initial studies used 7Hz stimulation of the vermis, which imposes a 7Hz (theta) oscillation on the hippocampus (Zeidler et al., 2020). Therefore, it was possible that the coordinated modulation of hippocampal interneurons required theta-range stimulation of the cerebellum. However, we saw similar response profiles when a single, 1s, light pulse was delivered to the cerebellum, indicating that a specific frequency of cerebellar modulation was not required to see the noted effects on hippocampal interneurons (**Fig. 6**).

This suggests that cerebellar stimulation (even without a 7Hz stimulation frequency) produces a response in hippocampal interneurons congruent with exploratory (locomotion or direct object investigation) behaviors. Anecdotally, previous work using electrical stimulation of the cerebellum in human epilepsy patients noted increased mental alertness and an increase in CSF norepinephrine (Cooper et al., 1976; Van Buren et al., 1978), suggesting possible involvement of the locus coeruleus. While the pathway(s) and intermediaries between the cerebellum and the hippocampus are still being elucidated, additional candidates include regions important for the control of theta (e.g. the medial septum and supramammillary region) and the nucleus incertus (Watson et al., 2019; Farrell et al., 2021; Froula et al., 2023). The nucleus incertus receives direct inputs from the cerebellum (Fujita et al., 2020), shows theta-modulated activity patterns (Ma et al., 2013), promotes behavioral arousal (Ma et al., 2017), and provides both direct and indirect (via collaterals to the medial septum) inhibition of somatostatin-expressing CA1 (and hilar) interneurons (Szőnyi et al., 2019). The supramammillary region also receives cerebellar input (Fujita et al., 2020), has direct and indirect (including via the medial septum) connections with the hippocampus (Vertes, 1992; Soussi et al., 2010; Li et al., 2020; Kesner et al., 2021), impacts hippocampal theta (Pedersen et al., 2017; Ruan et al., 2017; Ito et al., 2018; Billwiller et al., 2020) as well as processing of both novelty (Chen et al., 2020) and locomotion (Farrell et al., 2021), and is therefore potentially particularly relevant to our findings. Recent work in epileptic animals has also demonstrated that cerebellar outputs to the central lateral thalamus can play a key role in cerebellar modulation of the hippocampus, at least in the context of hippocampal seizure control (Streng et al., 2021). The central lateral thalamus is also believed to play a key role in arousal (Van der Werf et al., 2002; Schiff, 2008; Gummadavelli et al., 2015). Future work in healthy animals examining how cerebellar outputs can shape hippocampal activity, including that of CA1 interneurons, via the central lateral nucleus of the thalamus and via other intermediaries will be important to better understand cerebello-hippocampal communications and their functional roles.

There is a growing appreciation for the impact the cerebellum can have on hippocampal processing (Babayan et al., 2017; Lefort et al., 2019; Watson et al., 2019; Zeidler et al., 2020; Liu et al., 2022; Froula et al., 2023; Streng et al., 2023). The cerebellum can influence the firing of individual hippocampal neurons, as well as spatial navigation and the processing of spatial cues and landmarks, including the processing of objects in space. Our work indicates that these impacts extend to CA1 inhibitory interneurons, and that the cerebellum can help orchestrate CA1 circuits by activating and inhibiting CA1 interneuron populations in a coordinated manner.

## Acknowledgements

The authors would like to thank the entire K-M lab, especially past lab member Dr. Zachary Zeidler for imparting his knowledge for training in miniscope surgical, experimental, and analysis techniques, as well as Madison Smith for mouse colony management. Dr. Ezequiel Marron, manager of the University of Minnesota Viral Vector and Cloning Core, provided essential support by packaging the interneuron-specific vector from Addgene (initially developed by Dr. Gordon Fishell; plasmid #83899). We also thank Dr. Erin Lind for assistance with optogenetic equipment. Finally, we thank the UCLA Miniscope project for developing open-source materials for miniscope experiments and analysis.

## Supplemental Materials and Methods

**Supplemental Figure 1.**
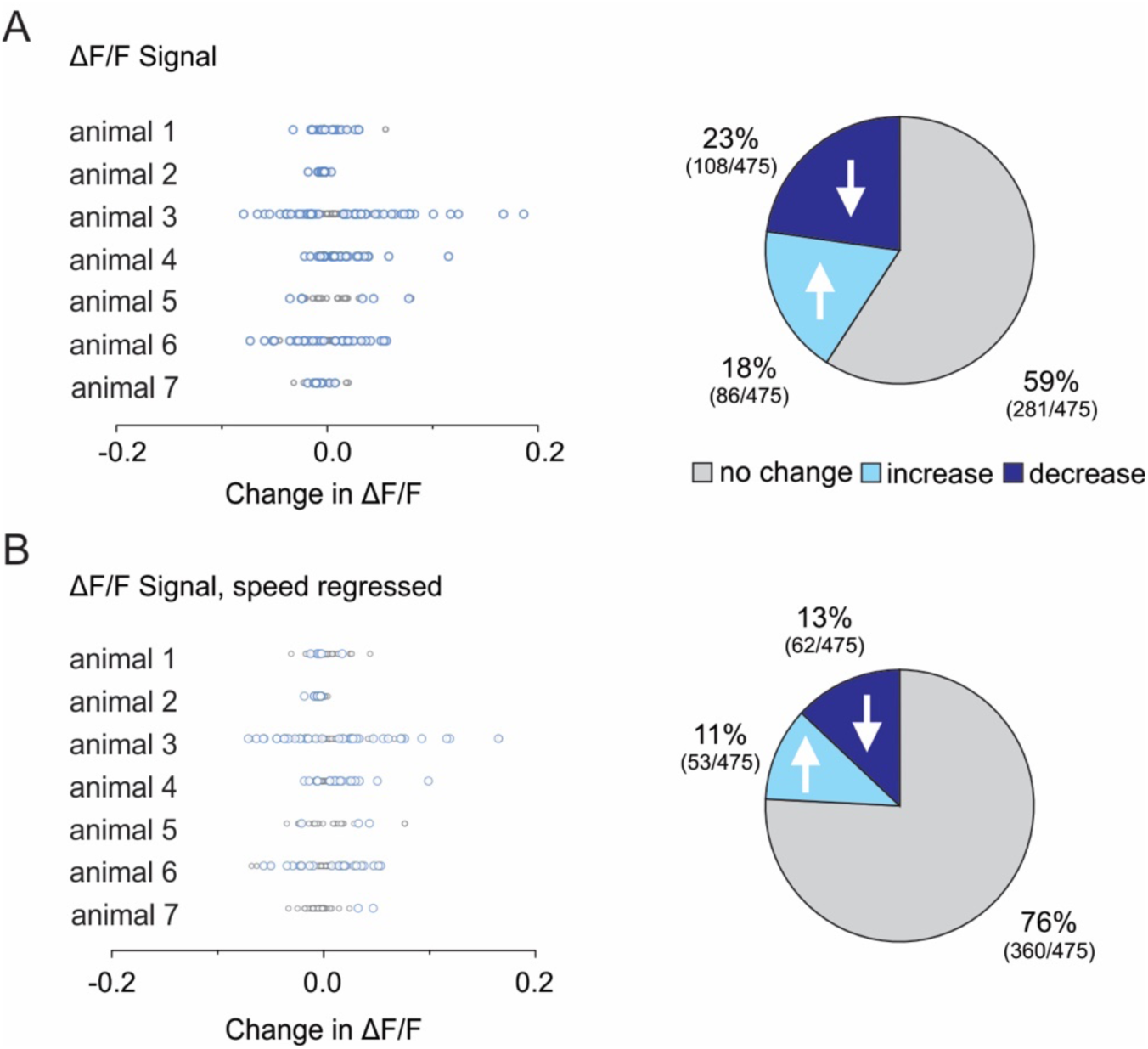
CA1 interneuron responses to cerebellar stimulation using ΔF/F signal and with speed regressed from ΔF/F signal. **A,** Responsiveness of CA1 interneurons to cerebellar stimulation using ΔF/F (in contrast to the deconvolved signal used for analyses represented in Figure 2). *Left:* Change in mean activity pre-stimulation compared to during stimulation. Each dot represents one cell. Blue: cells significantly modulated by light delivery. Gray: cells not significantly modulated by light. *Right:* Percentage and proportion of cells not responding to light (light gray), showing a significant decrease in activity (dark blue), or showing a significant increase in activity (light blue) in opsin-positive animals. **B,** Same as (A), but after removing the influence of speed (linear regression) from the calcium signal. Note that the ΔF/F signal rather than the deconvolved signal was used for this analysis due to the non-linear nature of the deconvolved signal (i.e. the deconvolved signal contains a large proportion of zeros). Residual calcium activity was compared, and a cell was considered significantly modulated by light beyond the influence of speed if the p-value for that cell was <0.05 (Wilcoxon Signed Rank test). Following speed correction, ∼24% of CA1 interneurons are significantly modulated by cerebellar stimulation, with approximately half of those activated and the other half inhibited during light delivery.

**Supplemental Figure 2.**
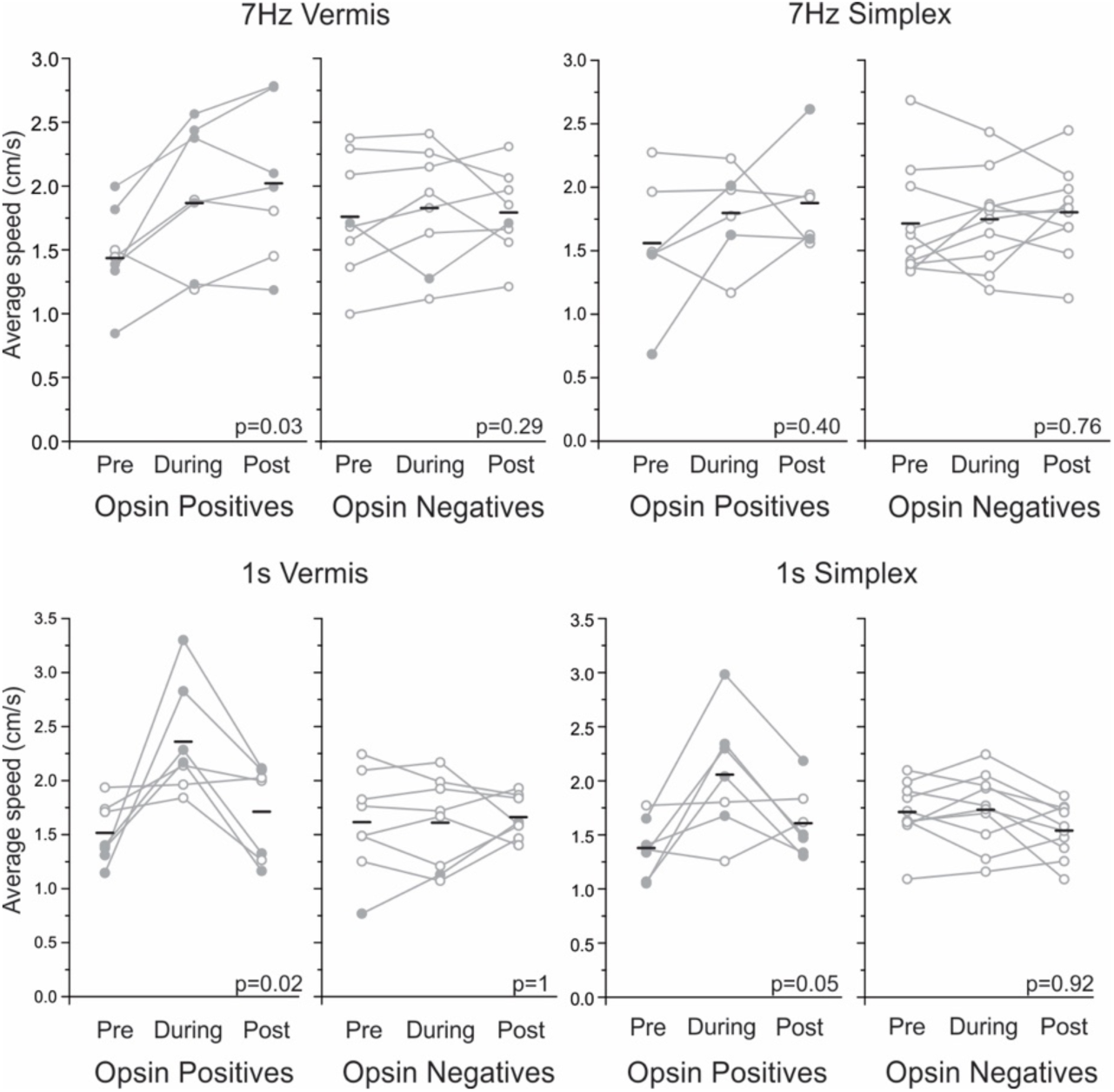
Locomotion around stimulation periods. Average speed by animal calculated across all stimulations (25-30 per session) by genotype and stimulation paradigm for the 3 s before stimulation (Pre), 3 s following the onset of stimulation (During), and 3 s post stimulation (Post). Filled circles represent animals with a significant change in activity from Pre to During (Wilcoxon Signed Rank Test), indicating a potential effect of cerebellar stimulation on locomotion. Bars represent means across animals. “Pre stimulation” locomotion and “during stimulation” locomotion were compared by genotypes in each stimulation paradigm (Wilcoxon) and are displayed in the bottom right corner of each graph.

**Supplemental Figure 3.**
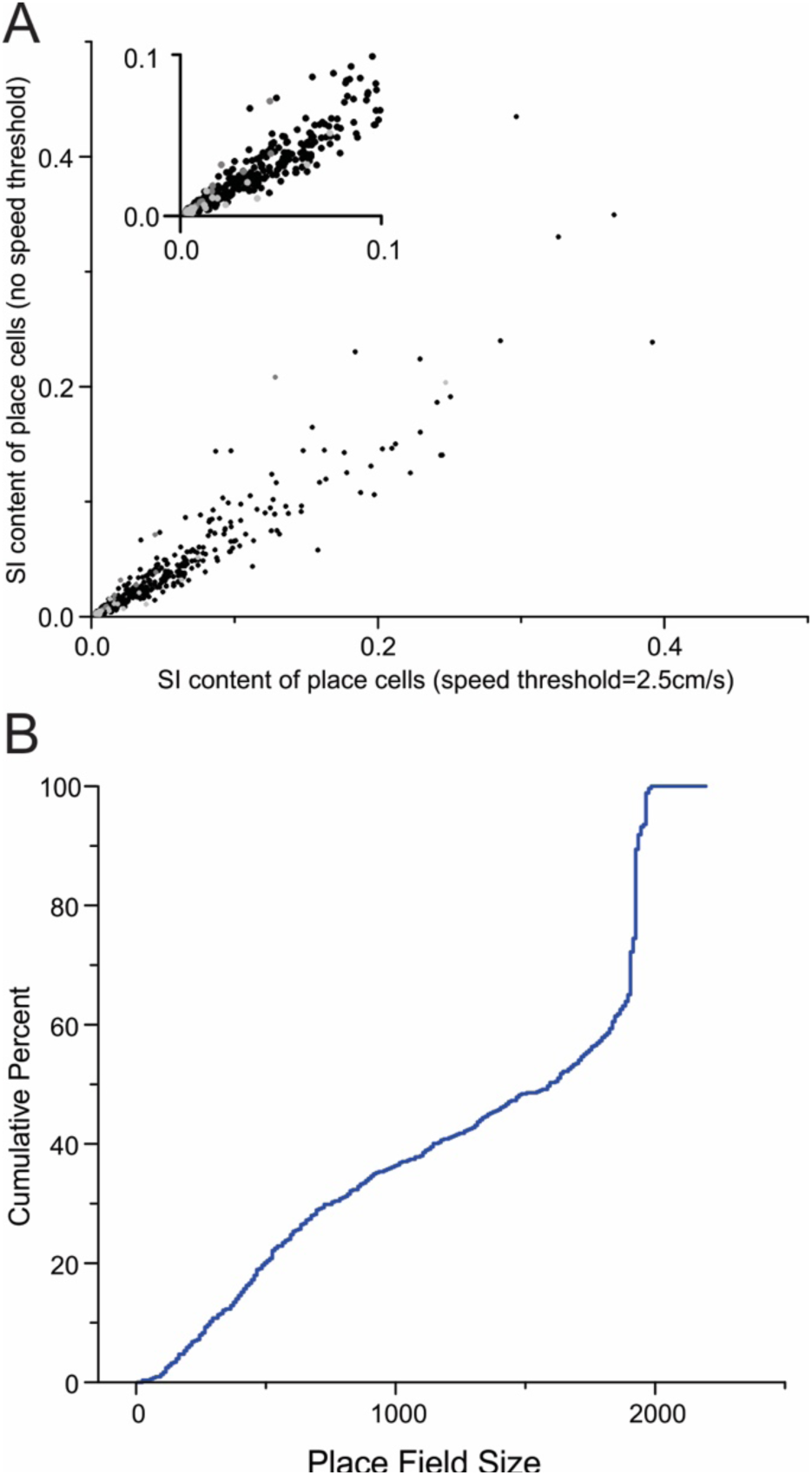
Hippocampal interneurons exhibit a range of spatial information content scores and place field sizes. **A,** Spatial information (SI) content scores of place cells in opsin negative animals using only fast epochs (i.e. animal traveling above 2.5cm/s; x-axis) or all speeds (y-axis). Only cells with SI scores from 0 to 0.5 are shown. Black dots: cells that were place cells in either condition; Dark gray dots: cells that were place cells only when including all speeds; Light gray dots: cells that were place cells only when using fast epochs. Inset: SI scores from 0 to 0.1. Note many place cells have low spatial information content but some interneurons also have higher SI scores, indicative of more restrictive spatial tuning. Note also that restricting analysis to periods of locomotion typically resulted in stronger SI scores. **B,** Cumulative distribution of place field sizes (units: cm^2^) in opsin negative animals. Note that place field sizes in interneurons are frequently quite large (compatible with previous reports showing broad spatial tuning in interneurons) but can also be specific to smaller portions of the arena.

**Supplemental Figure 4.**
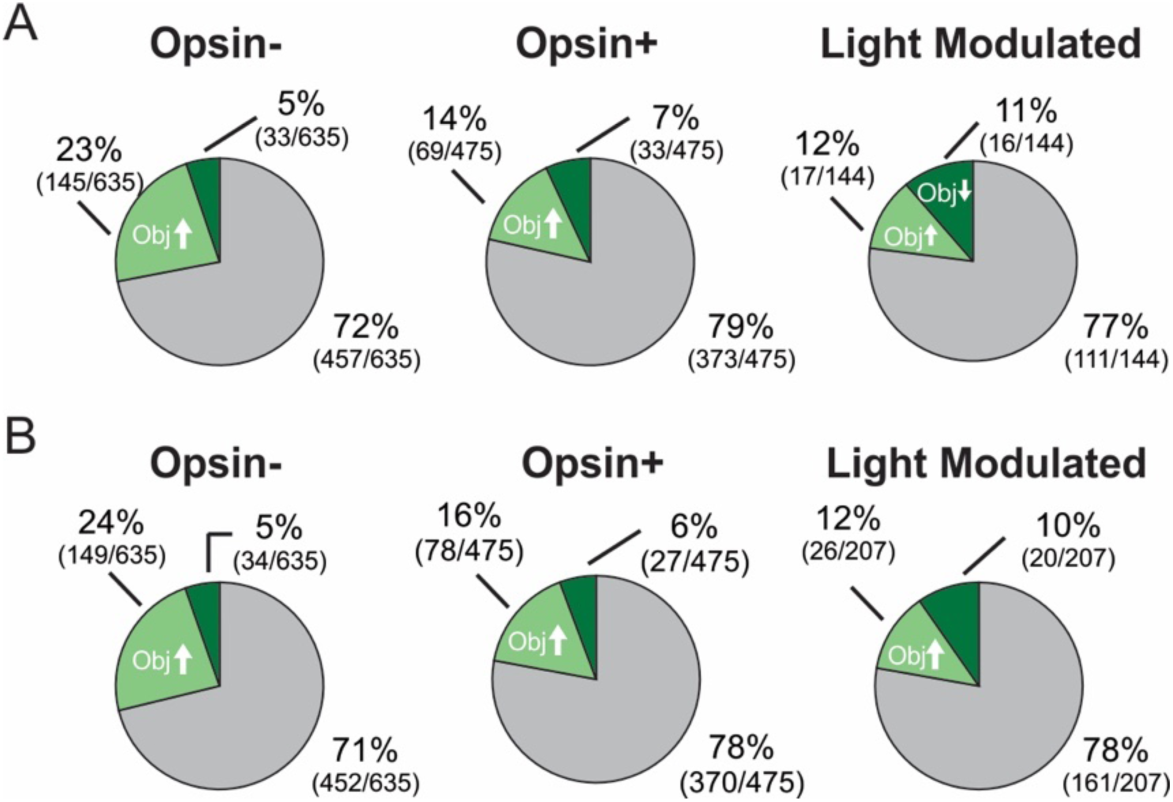
Speed correction does not meaningfully impact proportions of object-responsive CA1 interneurons. **A**, Number of Obj↑, Obj↓, and non-object-investigation modulated cells across all cells in opsin-negative animals (left), opsin-positive animals (middle), and for just light-modulated cells in opsin-positive animals (right), after speed correction. The correlation between speed and calcium activity was calculated for periods around object investigations – specifically the 7 s peri-investigation time period: the pre-investigation window (3 s baseline + 1 s grace period) and the start of the investigation window (3 s starting at the onset of investigation). Residual calcium activity was compared, and a cell was considered significantly modulated by object investigations beyond the direct influence of speed changes if the p-value for that cell was <0.05 (Wilcoxon Signed Rank test). Cells that increase their activity with object exploration are represented in light green, and cells that decrease their activity with object exploration are in dark green. **B,** Data repeated from **Fig. 3E and 3G** for reference, showing proportions without speed correction. Note that the proportions of object-responsive cells are largely unchanged when the influence of speed is removed. For all pie charts, the numbers in parentheses indicate the number of cells. Data drawn from 7 opsin-positive animals and 8 opsin-negative animals.

**Supplemental Figure 5.**
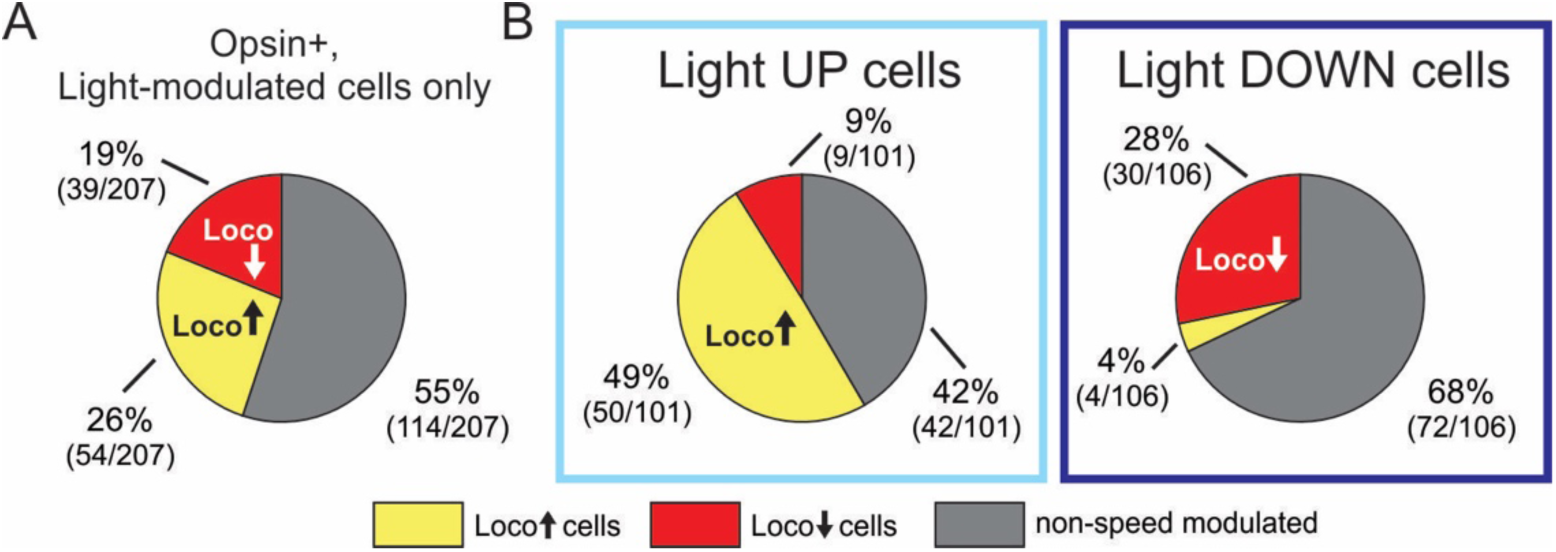
Categorization of light-modulated interneurons in CA1 using speedups. **A,** Proportions of Loco↑ (yellow), Loco↓ (red), and non-speed modulated (gray) cells around speedups for light-modulated cells from opsin-positive animals. **B,** As in A, but for Light↑ (left) and Light↓ (right) subpopulations. Note that most Light↑ cells that are locomotion modulated are Loco↑ cells (top), while most Light↓ cells that are locomotion modulated are Loco↓ cells (bottom), as was seen when instead using slowdowns for categorization (see Fig. 4). Data drawn from 7 opsin-positive animals.

## Supplemental Tables

**Supplemental Table 1:** Similar distributions found regardless of stimulation frequency. For functional categorizations using object exploration and slowdowns, the breakdown of Obj↑, Obj↓, and non-object modulated or Loco↑, Loco↓, and non-locomotion modulated were compared across stimulation frequencies. Number of animals represented for each comparison is included in parentheses under the “Behavior Category” description. Comparisons were made across all light modulated cells or by direction of light modulation (i.e. Light↑ or Light↓), and specific panels that were compared from Figure 6 are listed in the “Figure 6 Panels” column. Distributions are similar across different stimulation paradigms (note that the Bonferroni corrected alpha given multiple comparisons = 0.0042; however, none are significant even with an uncorrected alpha of 0.05), suggesting that a specific cerebellar stimulation paradigm is not necessary to achieve bidirectional, coordinated modulation of CA1 interneurons according to their functional roles.

| Comparison | Behavior Category | Cell category | Figure 6 Panels | Chi-squared P-value |
| --- | --- | --- | --- | --- |
| Vermis: 7Hz vs 1s | Object Exploration<br>(7Hz N=7; 1s N=4) | All Light-modulated cells | Ai vs Bi | 0.15 |
|  |  | Light Up cells only | Aii vs Bii | 0.16 |
|  |  | Light Down cells only | Aiii vs Biii | 0.67 |
|  | Locomotion via slowdowns<br>(7Hz N=6; 1s N=7) | All Light-modulated cells | Ei vs Fi | 0.18 |
|  |  | Light Up cells only | Eii vs Fii | 0.42 |
|  |  | Light Down cells only | Eiii vs Fiii | 0.78 |
| Simplex: 7Hz vs 1s | Object Exploration<br>(7Hz N=5; 1s N=7) | All Light-modulated cells | Ci vs Di | 0.23 |
|  |  | Light Up cells only | Cii vs Dii | 0.57 |
|  |  | Light Down cells only | Ciii vs Diii | 0.30 |
|  | Locomotion via slowdowns<br>(7Hz N=6; 1s N=7) | All Light-modulated cells | Gi vs Hi | 0.42 |
|  |  | Light Up cells only | Gii vs Hii | 0.06 |
|  |  | Light Down cells only | Giii vs Hiii | 0.51 |

